# The control of prickle formation in *Rubus*

**DOI:** 10.64898/2025.12.22.695586

**Authors:** Brian St. Aubin, Tom Poorten, Andrew Fister, Cherie Ochsenfeld, Joel Reiner, Allie Sandra Castillo, Rishi Aryal, Tomáš Brůna, Olga Dudchenko, Daniel James Sargent, Daniel Mead, Matteo Buti, Alexander Silva, Melanie Pham, David Weisz, Nahla Bassil, Hudson Ashrafi, Erez Lieberman Aiden, Nat Graham, Deepika Chauhan, Eric Dean, Warner Lowry, Lauren Redpath, Pradeep Marri, Shai Lawit, Gina Pham, Margaret Worthington, Brian CW Crawford

## Abstract

Prickles on blackberry and raspberry canes make pruning, harvesting, and handling more difficult and can increase labor costs for growers. The trait has been challenging to improve in these clonal crops because it is recessive and linked to undesirable agronomic traits. In blackberry and red raspberry, breeding programs have used recessive mutants at the *S* locus to generate prickleless cultivars for the last century. In this study, we identified independent loss-of-function mutations in a WUSCHEL-LIKE HOMEOBOX transcription factor, *WOX1*, as the genetic basis of the prickleless *S* locus in both blackberry and red raspberry. We mapped the *S* locus using integrated genome-wide association, bulked segregant analysis, and identity-by-descent analyses informed by breeding pedigrees. Additionally, we generated a genome sequence from Luther Burbank’s prickleless blackberry variety Burbank Thornless that contained an additional allele of *WOX1*. To verify the gene’s role, we used gene editing to knock out *WOX1* in an elite prickled commercial blackberry line. All edited plants were prickleless and lacked glandular trichomes, confirming that *WOX1* controls a joint developmental pathway. Other plant traits were unchanged, indicating *WOX1* is a specific and safe target for improvement. Gene editing can enable breeders to remove prickles directly from elite varieties, reducing the need for extensive breeding cycles and delivering safer, easier-to-harvest cultivars to growers.

## Introduction

In *Rubus* species like blackberry and red raspberry, prickles serve as a key defensive feature that protects the plant from herbivores and other threats. These sharp outgrowths, which arise from the non-vascular tissue, act as a physical barrier, deterring animals from feeding on or damaging the plant’s stems and leaves (Table 1). In addition to providing protection from grazing animals, prickles can also reduce damage from mechanical stress, such as wind or contact with other plants. While prickles are beneficial for the plant’s survival in the wild, they pose challenges in agricultural settings, particularly during harvesting and pruning, when they can hinder manual handling and increase labor efforts (Darrow, 1928; Jennings, 1986; Jennings, 1988). Because of this, breeding efforts in *Rubus* crops, collectively known as caneberries, often focus on developing prickle-free cultivars to improve efficiency and ease of cultivation (Finn and Clark, 2011).

**Table 1.**
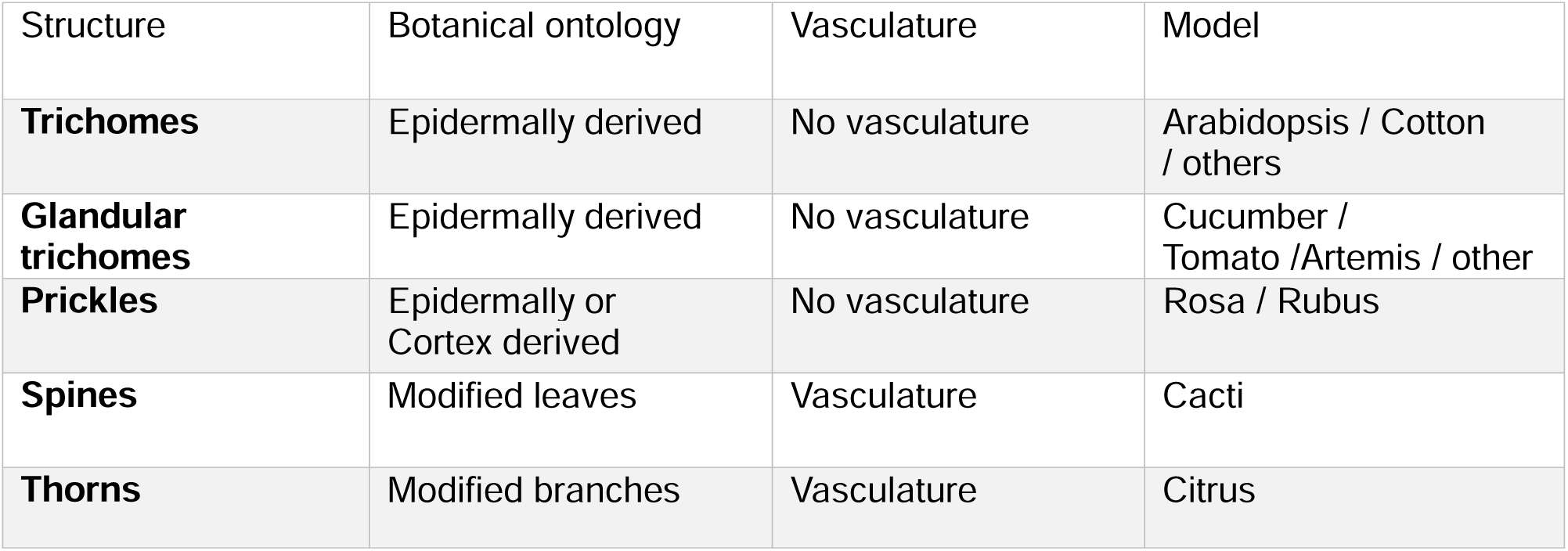
Differences in epidermal outgrowths.

| Structure | Botanical ontology | Vasculature | Model |
| --- | --- | --- | --- |
| **Trichomes** | Epidermally derived | No vasculature | Arabidopsis / Cotton / others |
| **Glandular trichomes** | Epidermally derived | No vasculature | Cucumber / Tomato /Artemis / other |
| **Prickles** | Epidermally or Cortex derived | No vasculature | Rosa / Rubus |
| **Spines** | Modified leaves | Vasculature | Cacti |
| **Thorns** | Modified branches | Vasculature | Citrus |

In *Rubus* species, both glandular trichomes and prickles are epidermal outgrowths that serve defensive roles, though they differ in structure and function (Khadgi and Weber, 2020a). Glandular trichomes are typically smaller, hair-like structures with secretory cells at their tips (distal heads) that produce compounds such as oils, terpenoids, or phenolics, which deter herbivores and protect against pathogens (Werker, 2000). In contrast, prickles are larger, sharp outgrowths that physically defend the plant by preventing herbivores or other animals from feeding on the stems or leaves. While glandular trichomes primarily offer chemical defense, prickles provide mechanical protection. Interestingly, both structures develop from the epidermis, and the genetic mechanisms underlying their formation may overlap. Studies in *Rubus* suggest that similar regulatory pathways could influence the development of both glandular trichomes and prickles, highlighting a potential genetic connection between chemical and physical defenses in the plant (Coyner et al., 2005). Understanding this relationship can aid in breeding efforts, especially for prickle-free cultivars, without compromising the plant’s natural defenses provided by glandular trichomes.

The genus *Rubus*, which is part of the Rosaceae family, is an ideal model for investigating the mechanisms behind prickle formation and development. The concept of prickleless canes in red raspberries (*Rubus idaeus*) was first documented in 1629 by John Parkinson in *Paradisi in sole paradisus terrestris* (Parkinson, 1629; Jennings, 1988) . Breeding for prickleless (also referred to as thornless) plants in caneberries has a long history that is driven by the need to improve harvesting efficiency and reduce labor costs associated with handling prickled plants. The quest for prickleless caneberry cultivars began in the early 20th century, with efforts focused on identifying naturally occurring prickleless mutations and incorporating them into breeding programs. In raspberries, the development of prickleless cultivars gained momentum with the discovery of the allele ‘spineless’ (‘*s^ri^*’, *R. idaeus*) in prickleless types of the progeny of the Scottish variety Burnetholm (Lewis, 1939). Subsequent breeding using this ‘*s*’ allele led to the creation of all the modern prickleless red raspberry varieties like Glen Ample and Joan J. (Jennings, 1986).

Prickleless blackberry (*Rubus* subgenus *rubus*) cultivars have been developed from multiple heritable and non-heritable mutations (Clark et al., 2007). Many early prickleless cultivars, including Thornless Evergreen, Thornless Loganberry, and Thornless Youngberry, were found to be chimeral, with the prickleless alleles only present in the non-heritable epidermal (L1) layer, complicating their use in breeding. A dominant, heritable source of prickleless canes was discovered in the octoploid cultivar Austin Thornless, but its use in modern breeding programs has been limited by its partial sterility, trailing habit, and presence of prickles in the basal 30 cm of canes. The recessive ’*s*’ allele (the ‘*s^ru^*’, *R. ulmifolius* allele) is by far the most used source of the prickleless trait in blackberry. The *s^ru^* allele is derived from the diploid *Rubus ulmifolius* var. *inermis*, also known as *Rubus rusticanus* var. *inermis*. Breeders generated a prickled tetraploid *R. ulmifolius* var. *inermis* cultivar, John Innes, carrying the recessive ‘*s*’ allele, from an unreduced germ cell (Crane and Darlington, 1927). The prickleless blackberry cultivar Merton Thornless was developed in the 1930s from a backcross of a prickleless F_2_ seedling to blackberry cultivar John Innes, and the *s^ru^* allele then served as a foundation for modern prickleless blackberry breeding (Crane and Darlington, 1927; Coyner et al., 2005). Luther Burbank’s most famous prickleless blackberry was a diploid variety of *R. ulmifolius* known as Burbank Thornless, which he promoted as an easier-to-handle alternative to traditional prickled varieties(Butterfield, 1928). Subsequent breeding advancements using Merton Thornless led to popular thornless blackberry varieties such as Navaho, Apache, Triple Crown, and Chester, all of which offer ease of cultivation and improved fruit quality (Coyner et al., 2005). These prickleless cultivars have become valuable in commercial production, significantly reducing the challenges associated with harvesting and field management while maintaining high fruit quality.

The *s^ru^* gene responsible for prickleless canes in blackberry was determined to be allelic with the *s^ri^* gene in raspberry(Jennings, 1986). Both *s^ru^*and *s^ri^* have been mapped to overlapping regions on *Rubus* chromosome 4 (Castro et al., 2013; Khadgi and Weber, 2020b; Johns et al., 2025) . There are a few limitations to using the *s^ru^* gene for achieving prickleless canes in blackberry breeding including recessive inheritance and associations with undesirable plant and berry traits. Modern fresh-market blackberry cultivars are tetraploids with multisomic inheritance.

Therefore, four copies of the recessive *s^ru^* allele are required for a plant to develop the prickleless phenotype. Early breeders using the *s^ru^* allele conducted self-pollination, sib-crossing, and backcrossing to develop prickleless plants, all of which contributed to inbreeding and loss of diversity around the *s^ru^* locus. This loss of diversity is still evident in modern breeding programs; an extensive linkage disequilibrium block was identified around the *s^ru^*locus in an association panel of fresh-market blackberry cultivars and breeding selections composed of 38 prickled and 336 prickleless genotypes (Johns et al., 2025) . Merton Thornless has several shortcomings as a cultivar, including susceptibility to freeze damage, trailing growth habit, late harvest season, large seed size, and high fruit acidity. Many of these traits appear to be in linkage with the *s^ru^* locus, and breaking up deleterious linkages has been challenging, particularly considering the recessive inheritance of the *s^ru^* allele and extensive linkage disequilibrium around the prickleless locus (Clark, 2005) .

Gene editing offers a solution to the problems associated with using the *s^ru^* allele in breeding programs by allowing precise genetic modifications, eliminating unwanted traits without affecting favorable ones (Moore, 1984) . Using this technology, breeders could modify the prickled allele(s) at the *s^ru^* locus in an elite prickled variety with exceptional fruit quality and agronomic traits without the time and expense of further crossing and the complications of potential inbreeding and linkage drag around the locus. This technology accelerates the breeding process and improves overall crop performance (Fister et al., 2024) .

In this paper, we identify mutations in a WUSCHEL-LIKE HOMEOBOX-type (WOX) transcription factor responsible for the prickleless phenotype in red raspberry and blackberry. We developed a new genome assembly and annotation for *R. ulmifolius* Burbank Thornless and mapped the ’s’ locus using a combination of genome-wide association studies (GWAS), bulked segregant analysis (BSA), and family-based identity-by-descent (IBD) mapping. We also developed a transformation protocol for blackberry and confirmed the target gene through gene editing, successfully introducing the prickleless phenotype into a commercial blackberry variety previously featuring prickles.

## Results

### Generating a diversity panel to understand the prickle trait in Rubus

We surveyed a diverse panel of 268 *Rubus* accessions, including 224 prickled and 44 prickleless accessions from eight subgenera, ensuring a broad representation of genetic and phenotypic variation for the prickle trait (Figure 1A; Supplemental Table 1). We observed a prickleless (smooth) phenotype in some *Rubus* subg. *rubus* and all *Rubus* subg. *cylactis*. Prickleless red raspberries (*Rubus* subg. *idaeobatus*) have been identified and bred, but none were included in the panel. Accessions were phenotyped with both unaided observation of the cane and with the use of a dissecting microscope (Figure 1A). Using qualitative visual evaluation, we distinguished four classes of phenotypes: plants with sharp, lignified prickles, plants with elongated, hairlike bristles, plants with both hairlike bristles and prickles, and smooth plants that lacked any elongated epidermal structure. When viewed with a dissecting microscope, we were able to better classify trichome types. Most plants had glandular trichomes, which varied in size and density. Some accessions had two unique glandular trichome structures: one short (<1mm) outgrowth, and one with an elongated stalk. With the microscope we were able to discern that the bristles observed by eye were these elongated glandular trichomes. All accessions had simple trichomes, which also varied in density among accessions. These typically laid flat along the cane, which sometimes made them difficult to observe without a microscope. In our study, we observed a Cramer’s V association statistic of 0.47 indicating a moderate correlation between the coded values for glandular trichomes and the type of prickles across the *Rubus* diversity panel. Varieties that exhibited prickles had glandular trichomes, suggesting a possible linkage or shared developmental pathway between these two epidermal structures (Figure 1B). Additionally, a subset of subgenus *Rubus* varieties that were prickle-free lacked glandular trichomes entirely, which reinforced the idea that the absence of prickles may be genetically or developmentally associated with the absence of glandular trichomes. This pattern suggests a potential genetic basis underlying the co-occurrence of these traits, which could be crucial for understanding the mechanisms that regulate epidermal outgrowths in *Rubus* species.

**Figure 1.**
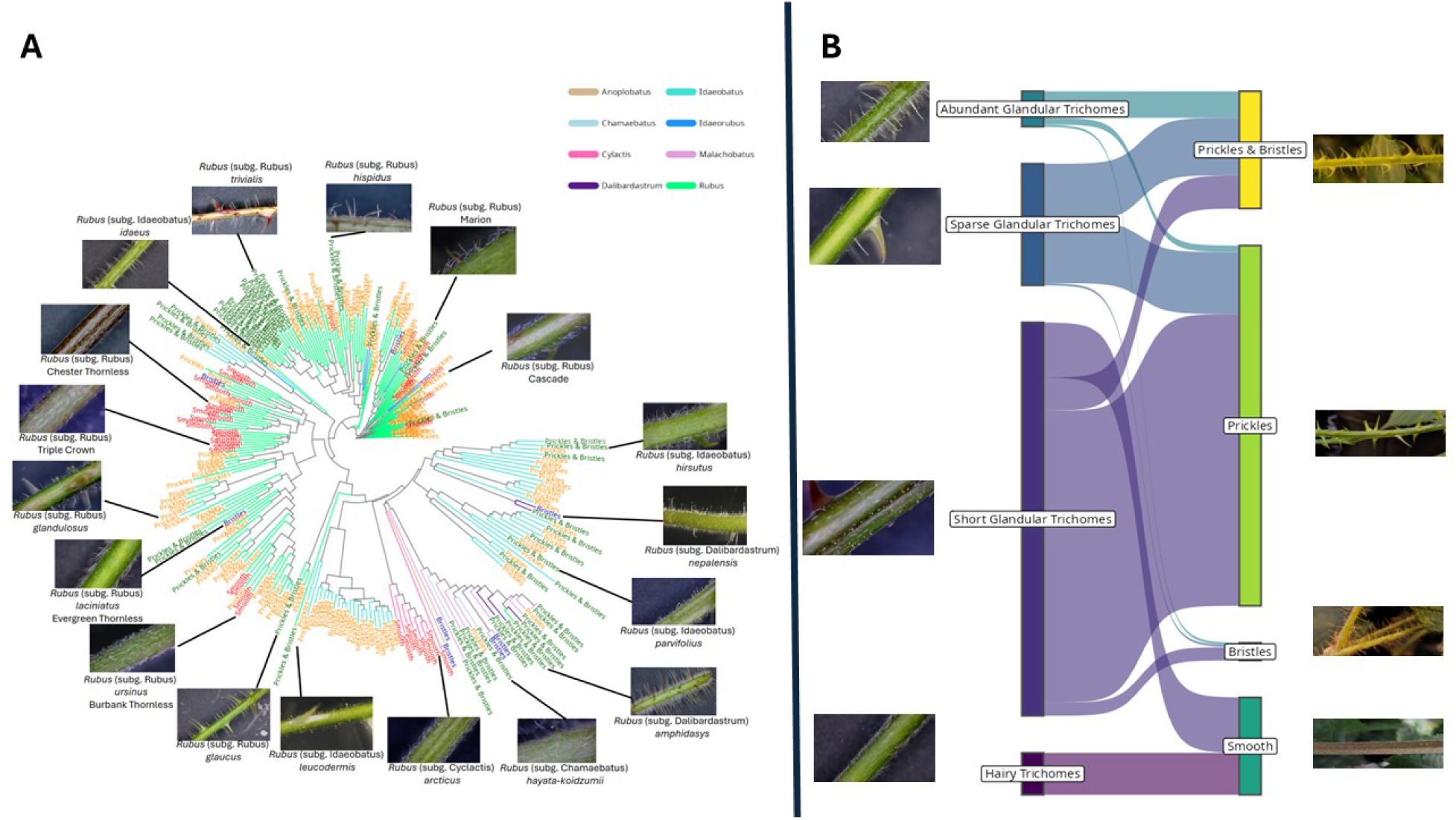
The morphological and genetic variation within the *Rubus* population illustrating the comparative morphology of trichomes, glandular trichomes and prickles. Panel A includes representative photographs showing the diversity of epidermal outgrowths on *Rubus* canes, as observed by eye. Panel B includes representative images of the diversity of trichome, glandular trichome, and prickle presentations as viewed under a microscope. Colored arrows indicate different epidermal structures: yellow = simple trichomes, red = short glandular trichomes, green = elongated glandular trichomes, blue = prickles. The scoring system indicates a wide range of variability within the panel, with a subset of individuals exhibiting both prickle-free and glandular trichome traits.

The inheritance of the “spineless” gene in red raspberry and blackberry has been well-documented through traditional breeding studies. In red raspberry, the *s^ri^* allele was successfully incorporated into various commercial cultivars. Within the diversity panel we identified both prickled and prickleless related red raspberry lines that were descended from crosses using the *s^ri^* allele (Supplemental Figure 1A). Similarly for blackberry, we identified samples within our diversity panel that contained the *s^ru^* allele from the original Merton Thornless accession (Supplemental Figure 1B). To aid the identification of the S locus we analyzed samples within our diversity panel that contained either of the *s^ru^*and *s^ri^* alleles (Supplemental Figure 1; Supplemental Table 2; Supplemental Table 3) through a combination of genome-wide association (GWAS), bulked segregant (BSA), and inheritance by descent (IBD) analyses.

### Developing a Burbank Thornless genome assembly and annotation

Like the progenitor of Merton Thornless, Burbank Thornless is also a prickleless diploid *R. ulmifolius,* but the providence of this line was unknown. Luther Burbank may or may not have obtained the same prickleless genotype used to introgress the *s^ru^* allele into modern blackberry breeding programs. To gain genomic insight into the basis of the prickleless trait, we generated a chromosome-scale assembly of Burbank Thornless using Pacific Biosciences long-reads scaffolded with high-throughput chromosome conformation capture (Hi-C). The final assembly comprised 342 Mb across 288 scaffolds, including seven chromosome-length scaffolds totaling 299 Mb (87% of the genome). Structural annotation predicted 29,708 protein-coding genes and 38,105 coding transcripts, with an average gene length of 3.4 kb (Table 3). Benchmarking with BUSCO identified 96.2% of conserved eudicot genes, and most loci were further supported by EST alignments, peptide homology, and high annotation confidence scores (C-scores, which reflect the strength of external evidence supporting each gene model). Functional annotation assigned putative functions to most transcripts across multiple databases and domain resources. Comparative analyses revealed strong collinearity with the *R. argutus* ‘Hillquist’ genome and with other published diploid *Rubus* genomes, reflecting conserved chromosome structure across the genus (Brůna et al., 2022) . Notably, the Burbank Thornless gene set is smaller but more completely annotated than those of other *Rubus* genomes, reducing the likelihood of fragmented or spurious predictions and thereby enhancing its utility for candidate gene discovery. Complete methods and results describing the genome assembly and annotation are provided in Supplemental File 1 and Supplemental Tables 4 and 5.

**Table 2.**
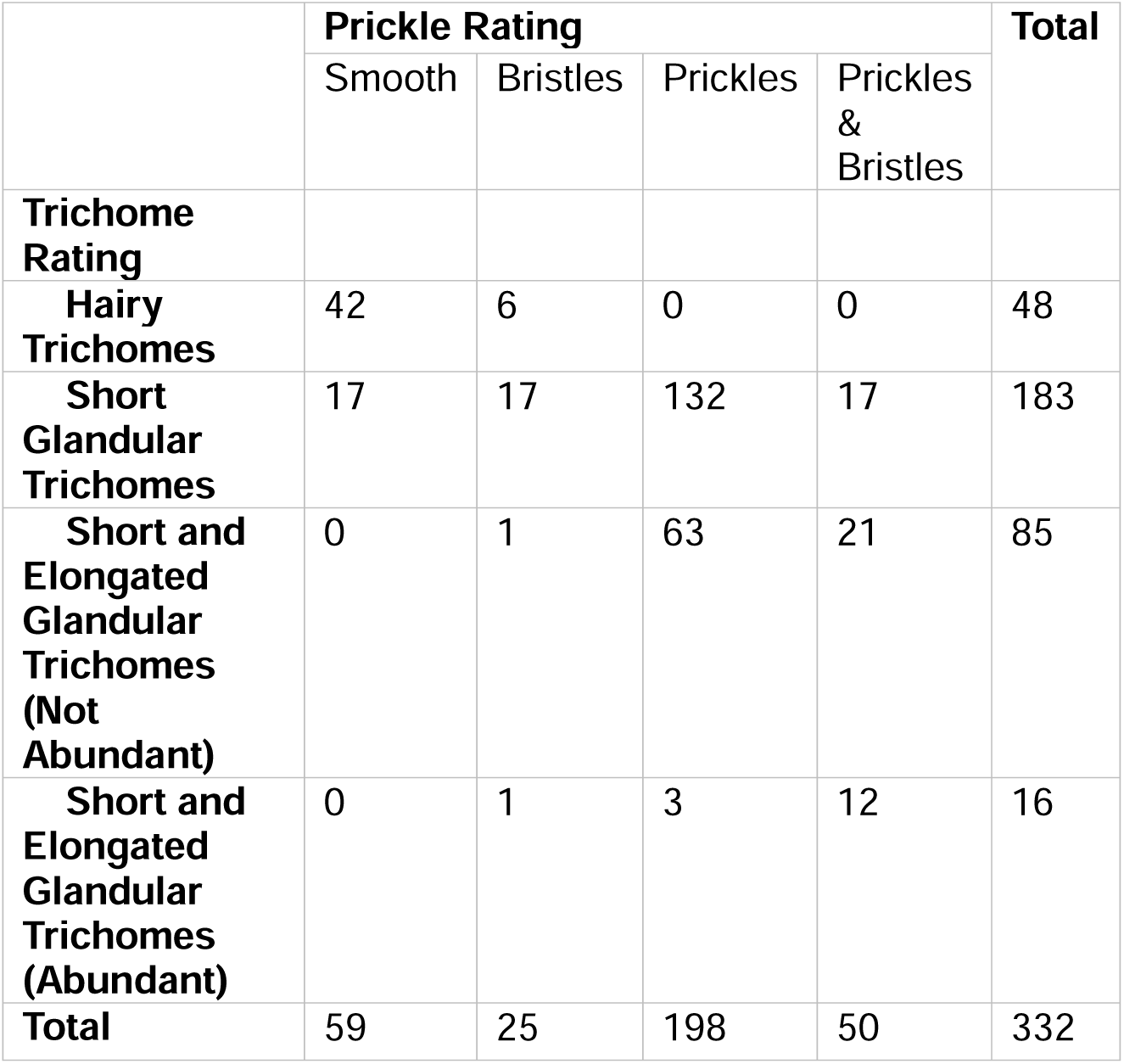
Prickle/trichome relationship in diversity panel.

|  | Prickle Rating |  |  |  | Total |
| --- | --- | --- | --- | --- | --- |
|  | Smooth | Bristles | Prickles | Prickles & Bristles |  |
| **Trichome Rating** |  |  |  |  |  |
| **Hairy Trichomes** | 42 | 6 | 0 | 0 | 48 |
| **Short Glandular Trichomes** | 17 | 17 | 132 | 17 | 183 |
| **Short and Elongated Glandular Trichomes (Not Abundant)** | 0 | 1 | 63 | 21 | 85 |
| **Short and Elongated Glandular Trichomes (Abundant)** | 0 | 1 | 3 | 12 | 16 |
| **Total** | 59 | 25 | 198 | 50 | 332 |

**Table 3.**
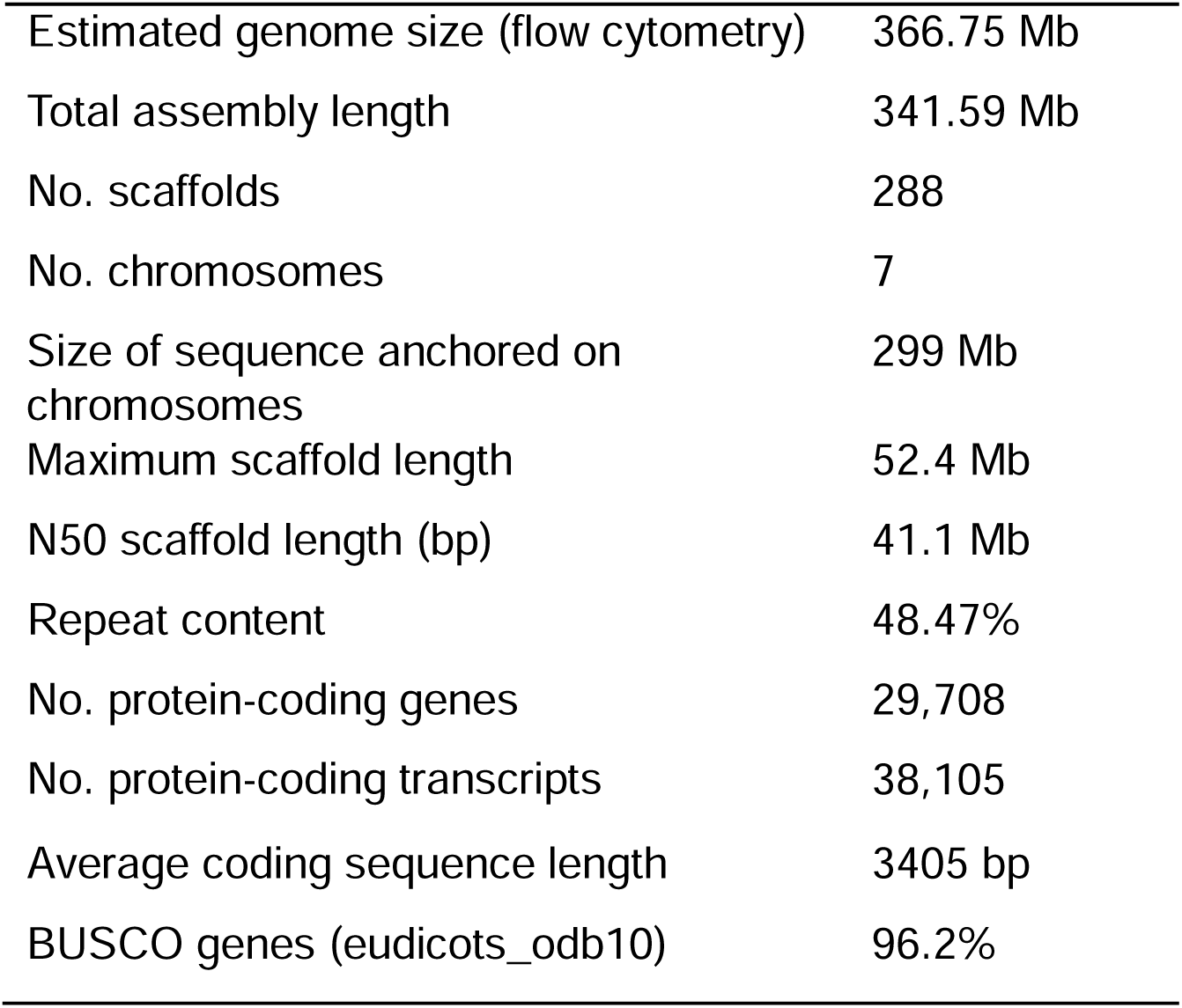
Summary statistics for the assembled *R. ulmifolius* cv. Burbank Thornless genome.

|  |  |
| --- | --- |
| Estimated genome size (flow cytometry) | 366.75 Mb |
| Total assembly length | 341.59 Mb |
| No. scaffolds | 288 |
| No. chromosomes | 7 |
| Size of sequence anchored on chromosomes | 299 Mb |
| Maximum scaffold length | 52.4 Mb |
| N50 scaffold length (bp) | 41.1 Mb |
| Repeat content | 48.47% |
| No. protein-coding genes | 29,708 |
| No. protein-coding transcripts | 38,105 |
| Average coding sequence length | 3405 bp |
| BUSCO genes (eudicots_odb10) | 96.2% |

### Fine mapping the prickleless causal mutation at the *S* locus

Three different mapping strategies were applied to resolve the causal locus. First, GWAS was used to find the locus associated with prickles on the whole diversity panel and on subsets of the diversity panel based on prior knowledge of the potential source of the causal allele. We also generated a segregating population for bulked segregant analysis in blackberry. Finally, we used pedigrees to inform an identity-by-descent (IBD) analysis to further refine the causal locus and manually inspected sequence variation in red raspberry and blackberry lines to look for recessive mutations in annotated genes.

We used a SNP-based GWAS to investigate the genetic basis of prickle development in a *Rubus* population of 43 accessions using the software package GWASpoly (Lemane et al., 2022; Voichek and Weigel, 2020). We scanned for associations between the presence of a homozygous mutant allele and the presence of the prickleless phenotype and were able to identify regions of the genome potentially linked to the *S* gene. We identified a region significantly associated with the prickleless trait at the end of chromosome Ra04 between 33,000,000 bp and the end of chromosome at 36,116,165 on the *R. argutus* genome (35,312,865 to 38,546,343 bp in *R. ulmifolius*) (Figure 2A, B; Supplemental Figure 2). This region overlaps with another region significantly associated with the prickleless trait that was recently identified in a GWAS panel of 374 fresh-market blackberry genotypes from 30.48 to 36.04 Mb on chromosome Ra04 (Johns et al., 2025) .

**Figure 2.**
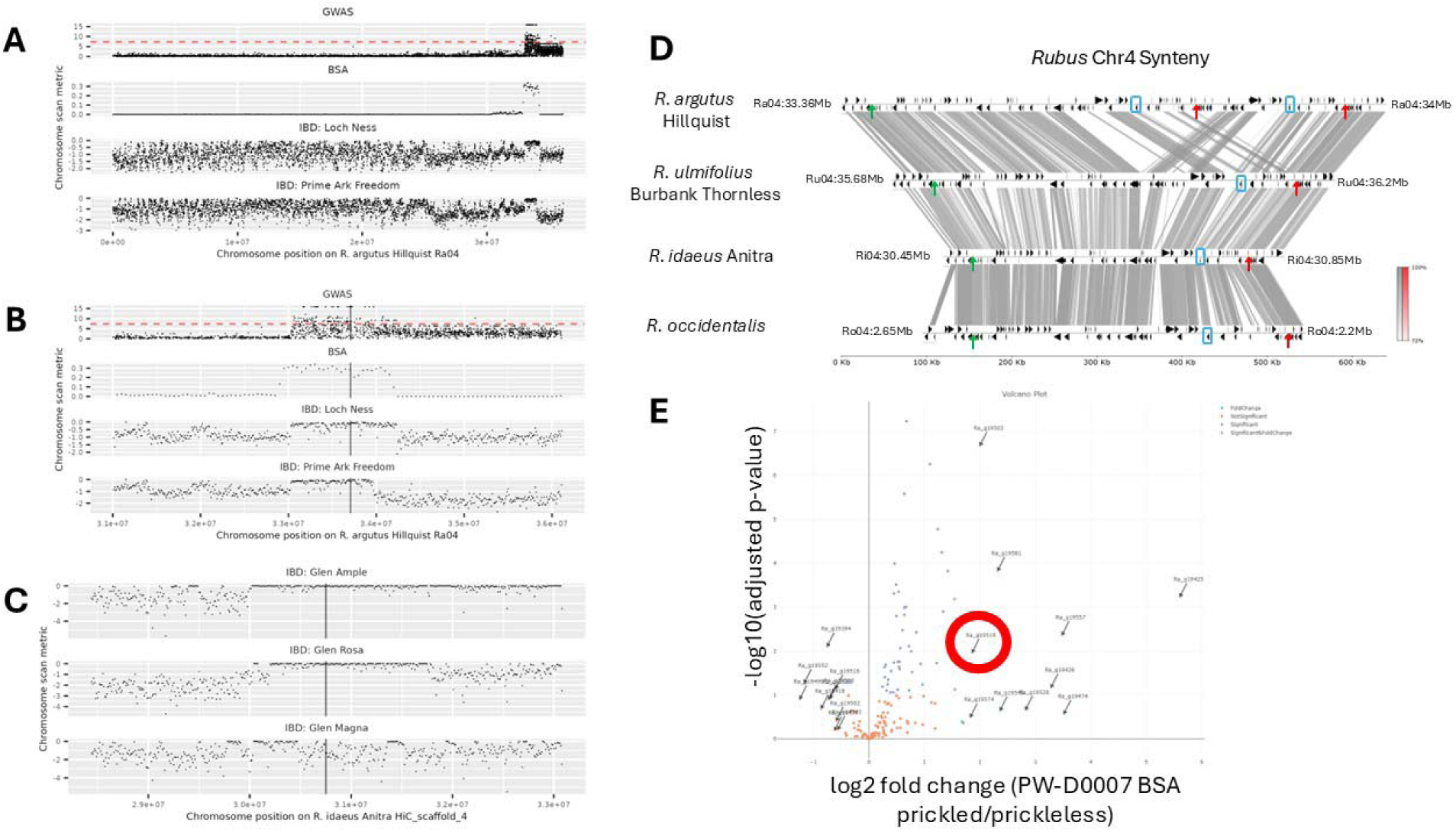
Fine mapping of *s* allele on *Rubus* chromosome 4. (A) *Rubus* chromosome 4 SNP density plots for the GWAS, BSA, and example individual lines mapped to the *R.argutus* cv. Hillquist genome. (B) Zoomed in view of 5Mbp region centered on pricklelessness-associated region on chromosome 4 in *R. argutus* Hillquist genome. (C) Zoomed in view of 5Mbp region centered on syntenic pricklelessness-associated region on chromosome 4 in *R. idaeus* Anitra (red raspberry) genome. GWAS panels shows –log10(p) on y axis, dashed red line shows standard significance threshold *p*=5x10^-8^. BSA, bulk segregant analysis, panels show number of filtered variants per 100kb window with a 50kb step size on the y-axis. IBD, identity-by-descent, panels show a metric to identify regions of excess homozygosity on y-axis (-[# heterozygous sites per 100kb window / median of # heterozygous sites per 100kb window]). (D) Synteny plot showing homologous pricklelessness-associated region in four Rubus genomes. Grey bands show mummer alignments (>=100bp and >=70% identity). Black arrows correspond to gene locations. Green arrows show syntenic positions of left border and red arrows show syntenic positions of right border. The Blue box shows locations of syntenic *WOX1* gene models. The region between left- and right-boundary in *R. argutus* is Ra04:33,393,466-33,774,434 (Ra04:33,951,000 is right-border when including tandem duplication misassembly); in *R. ulmifolius* is Ru04:35,726,940-36,153,835; in *R. idaeus* is Ri04:30,482,969-30,807,700; in *R. occidentalis* is Ro04:2,224,711-2,591,926. For visual clarity of synteny, the *R. occidentalis* chromosome 4 was reverse-complemented prior to alignment and plot generation. The Grey and Red scale bars show alignment sequence similarity for forward and reverse alignments, respectively. (E) Gene expression volcano plot for genes in the IBD interval, PW-D0007 siblings segregating for prickles. The Red circle highlights *WOX1*.

To further refine the candidate region, we generated a segregating population by self-crossing PW-D0007, a line triplex for the *s* allele (*Ssss*). The resulting progeny displayed either the prickleless (32 individuals) or prickled phenotype (65 individuals). We used this population in a bulked segregant analysis. High-throughput sequencing was then performed on both groups, generating genome-wide sequence data representing the genetic makeup of the prickleless and prickled groups (Figure 2A, B).

Given the tetraploid nature of our blackberry genotypes, we accounted for the complexity of allele dosage, where individuals can have varying numbers of copies of a given allele (e.g., zero to four copies). To handle this complexity, we focused on identifying regions of the genome where allele frequencies significantly differed between the two bulks. By comparing the allele frequencies in the prickled versus prickleless bulks, we were able to pinpoint the genomic region enriched for prickleless alleles to a 1.3 Mb region on chromosome Ra04 between 32,950,000 and 34,250,000 base pairs (35,259,235 to 36,474,787 in *R. ulmifolius*) (Figure 2 A,B).

In our efforts to fine map the locus responsible for the *S* gene, we also utilized a family-based identity-by-descent (IBD) analysis. We selected a subset of prickleless and prickled individuals from blackberry and raspberry along with prickled black raspberry, focusing on individuals known to have the *s^ru^*or s*^ri^* alleles to maximize shared genomic regions inherited from common ancestors (Supplemental Figure 1A,B). Using genotype information from the GWAS resequencing data, we identified regions of extended homozygosity in individuals without prickles and in individuals with prickles. By comparing these shared homozygous regions across the prickleless population, we were able to identify genomic intervals potentially harboring the gene of interest. We used both red raspberry and blackberry alleles to narrow the region further. We compared SNPs in heterozygous and homozygous individuals in a 20 kb region near the borders of the locus (Figure 2 A,B; Table 4). We identified that the prickleless cultivar Prime-Ark® Freedom had a recombination event that narrowed the region of homozygosity on the left side to 33,286,753 base pairs on the *R. argutus*, cv. Hillquist chromosome Ra04 (35,349,585 to 36,170,216 in *R. ulmifolius*) (Table 4). The *R. argutus*, cv. Hillquist genome contains a 166kb duplication within the region, impacting the identification of the right border. To resolve this issue both the *R. occidentalis* v3 and *R. idaeus* cv. Anitra genomes were primarily leveraged to identify SNPs (VanBuren et al., 2016; Davik et al., 2022) . The right border was set at Ri04:30,807,700 which corresponds to Ra04:33,951,000 and 33,774,434 in the *R. argutus* cv. Hillquist genome due to the duplication (Figure 2D). This narrowed region contained 82-140 predicted annotated genes depending on the genome and annotation. In red raspberry, the prickleless Glen Ample line shared homozygosity within the previous identified region with the related prickled line Glen Magna. This identified SNPs that provided a discrete boundary within the region at position 30,482,969 on chromosome 4 in the *R. idaeus* cv. Anitra genome (corresponding with chromosome Ra04, position 33,393,466 in *R. argutus*, cv. Hillquist and 35,726,846 in *R. ulmifolius*) (Figure 2C). This reduced the region down to around 324 kb containing 62-115 gene models (Supplemental Table 6).

**Table 4.** Genetic intervals potentially containing the causal mutation for the prickleless phenotype identified through fine mapping.

| Source allele | Method | Change | Left/right border relative to Hillquist genome | *Rubus argutus*, cv. 'Hillquist' location (bp) | *Rubus ulmifolius* Burbank Thornless location (bp) | *Rubus occidentalis* ORUS 4115-3 location (bp) | *Rubus idaeus* cv. Anitra1611 location (bp) | Nearest gene (inside) |
| --- | --- | --- | --- | --- | --- | --- | --- | --- |
| Red raspberry | Family based analysis | A>T homozygous to heterozygous | Left | Ra04:33,393,466 | Ru04:35,726,846 | Ro04:2,591,816 | Ri04:30,482,969 | Ra_g19461, Ruulm.4G300300, Ro04_G00141, Ri04_g21576 |
| Blackberry | Family based analysis | A>G | Left | Ra04:33,286,753 | Ru04:35,604,782 | Ro04:2,714,466 | Ri04:30,373,221 | Ra_g19440, Ruulm.4G298200, Ro04_G00118, Ri04_g21556 |
| Blackberry | Family based analysis | T>C | Right | Ra04:33,951,000 and 33,775,434 | Ru04:36,151,812 | Ro04:2,223,785 | Ri04:30,807,700 | Ra_g19578 and Ra_g19536, Ruulm.4G307000, Ro04_G00205, Ri04_g21637 |

**Table 5:** Differential gene expression within interval.

| Gene_ID | start | stop | strand | Gene info | domain info | PW-D0007 prickled/prickless fold change |
| --- | --- | --- | --- | --- | --- | --- |
| **Ra_g19503** | 33,629,274 | 33,630,632 | - |  | Peptidase S10 | 2.00 |
| **Ra_g19519** | 33,705,382 | 33,707,440 | - | Indels in thornless genomes | Homeobox domain | 1.86 |
| **Ra_g19534** | 33,761,529 | 33,762,047 | - |  | C2 domain | 1.24 |
| **Ra_g19555** | 33,838,869 | 33,850,045 | + |  | Phosphoribulokinase/uridine kinase | 1.54 |
| **Ra_g19557** | 33,853,606 | 33,854,114 | - | gene model errors first, second, third exon. | SLC26A/SulP transporter | 3.48 |
| **Ra_g19577** | 33,943,104 | 33,943,622 | - |  | C2 domain | 1.22 |

### Expression analysis of candidate gene targets

To gain a better understanding of biological pathway differences in prickled and prickleless canes, we used transcriptome analyses to investigate gene expression differences in these two groups. This information aided in the evaluation of candidate genes from the locus. We generated transcriptome data from the PW-D0007 blackberry self-crossed population that was used for BSA, which included samples from plants segregating for the prickleless and prickled phenotypes. The dataset comprised meristem samples from growing canes with the intention of having part of the sample contain cells that are transitioning to prickles. We used non-bulked samples from 3 prickleless and 9 prickled plants. This approach aimed to identify differentially expressed genes linked to the presence or absence of prickles, with several genes also found within the IBD region. The results provided extra support for SNPs within genes associated with all *s* alleles.

Expression analysis revealed 2357 upregulated genes in the prickled samples and 1921 down regulated in the prickled samples compared to the prickleless samples (padj <0.05) (Supplemental Table 7). Within the region there were several genes that had higher expression in the prickled samples including Peptidase S10 Ra_g19503, Homeobox Ra_g19519, C2 domain Ra_g19534, Phosphoribulokinase/uridine kinase Ra_g19555, SLC26A/SulP transporter Ra_g19557, and C2 domain Ra_g19577 (Table 5; Figure 2E). No genes within the region had lower expression in the prickled samples compared to the prickleless samples.

### Identification of the *S* gene

To understand the potential causal genes within the region, we attempted to use computational genome annotation, which has become instrumental in exploring gene functions across the numerous new genomes. However, standard genome annotation poses challenges in making refinements based on new data or integrating valuable user input. Additionally, despite apparent synteny, sequence variations between *Rubus* species complicated alignment to a single reference genome, requiring a multi-genome approach.

To address these challenges, we inspected gene models within the target interval and compared them to gene expression data for red raspberry, black raspberry, and blackberry. This comparison was facilitated by aligning three separate genome browsers simultaneously allowing us to inspect SNPs from various whole genome sequencing efforts in the context of several genomes. Using the *R. argutus* cv. Hillquist, *R. occidentalis* v3 genome, and the *R. idaeus* cv.

Anitra genome from rosaceae.org as references, we confirmed comparable genomic regions. Using the pyGenomeViz package with BLAST alignments and default parameters (https://github.com/moshi4/pyGenomeViz), we aligned the following regions: Ru04:35300000:36300000 (+), Ri HiCscaffold_4:30206071-30307189 (+), Ro04:1900000-3050000 (-), and Ra04:33000000-34000000 (+) (Figure 2 D). While some inconsistencies were noted in gene models across these genomes, the coding sequences for most expressed genes appeared correctly annotated. After finalizing the interval via IBD analysis, we made manual adjustments to the gene models within this region to increase annotation accuracy.

Through this refined approach, we identified an 8bp insertion in the second exon of the Ra_g19519 (Ruulm.4G305500) gene within the prickleless haplotype in blackberry (Figure 3A, B). Ra_g19519 encodes a WUSCHEL-Related Homeobox (WOX) protein, which is vital for balancing stem cell maintenance and differentiation within meristems in plants. In Arabidopsis, members of the *WOX* gene family have overlapping roles in various meristem regions, where they contribute to the development of plant lateral organs (Haecker et al., 2004; Park et al., 2005; Wu et al., 2005; Breuninger et al., 2008; Shimizu et al., 2008; Nakata et al., 2012; Qian et al., 2021) (Figure 3G). Phylogenetic analysis aligned Ra_g19519 (and its black raspberry orthologue Ro04_G00187) with the Arabidopsis *WOX1* gene (At3g18010). For simplicity, we designated Ra_g19519 as *RuWOX1* in blackberry (Figure 3F).

**Figure 3.**
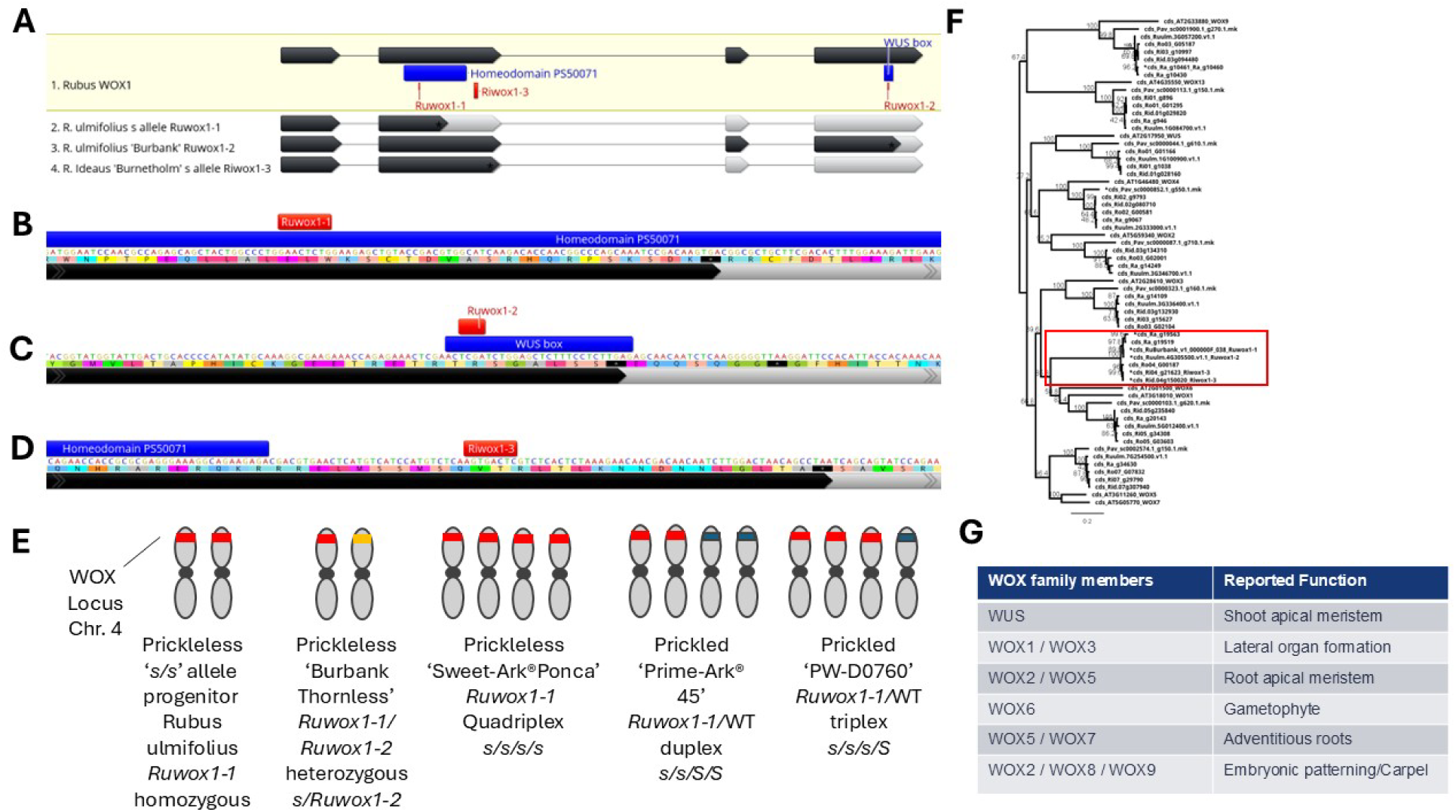
Mutant alleles and *WOX* gene phylogeny. (A) *Rubus* wild-type *WOX1* gene with dark grey showing CDS with regions labeled that encode for homeobox and WUS box protein domains. The *s* alleles are labeled and light grey showing truncation due to premature stop codons. (B-D) Close up of alleles with red indicating the insertions leading to premature stop codons. (E) Zygosity of blackberry *s* alleles in Blackberry lines. Red indicates *s* allele, orange indicates allele from Burnbank thornless and blue indicates wild-type *S* allele. (F) Phylogenomic tree of the conserved coding sequences (including indels) form closely related sequences to *WOX1*. *WOX13* and *WOX9* are included as an outgroup. The *WUS* sequence was not found in the *R. argutus* genome assembly and is substituted by the sequence from *R. ulmifolius*. The *WOX2* sequence was not found in the *R. ideaus* cv. Anitra genome assembly. Sequences with stop-codon inducing in-dels are marked with an *. (G) Arabidopsis *WOX* naming and the associated functions based on mutants and gene expression.

The 8bp insertion, termed *Ruwox1-1*, co-segregated with the prickleless (*s^ru^*) allele across all examined populations (Figure 3E). The insertion introduces a premature stop codon within the annotated homeodomain of *RuWOX1*, likely disrupting its function (Figure 3B). During IBD analysis of the blackberry *s^ru^* allele, we encountered difficulties in reconciling the prickle-free phenotype of the Burbank Thornless cultivar, which appeared heterozygous for the *Ruwox1-1* allele. Closer examination of the second Ra_g19519 allele in Burbank Thornless (unassembled contig 000000F_038) revealed two distinct haplotypes at the *S* locus, leading to the discovery of an additional allele, *Ruwox1-2*. *Ruwox1-2* carries a 4bp deletion within the WUS box, a conserved motif critical to the function of *WOX* family genes (Figure 3C). This finding indicates that Burbank Thornless has a compound heterozygous genotype at the *S* locus, with one copy of *Ruwox1-1* and one of *Ruwox1-2* (Figure 3E). Because the Burbank Thornless assembly is not phased, only one allele at the *S* locus was retained in the final reference sequence. By chance, the assembled haplotype corresponds to *Ruwox1-2*, the allele that was not introgressed into domesticated germplasm, while the alternative allele (*Ruwox1-1*) segregates with the prickleless trait in modern breeding populations. In contrast, our prickled commercial line, PW-D0760(Adams, 2022), was found to carry the *Ruwox1-1* allele in triplex, retaining one functional copy alongside three *s^ru^* alleles. Further analysis of Ri04_g21623, the red raspberry orthologue of Ra_g19519, identified an 8bp insertion just downstream of the homeodomain that introduces a premature stop codon before the last two exons (Figure 3D). We named this allele *Riwox1-3*, and it segregated with the prickleless (*s^ri^*) allele in red raspberry populations derived from the original Burnetholm cultivar. While evaluating phenotypic diversity across *Rubus* accessions, we observed that most *Cylactis* species exhibited a smooth phenotype. This observation prompted an investigation into potential *WOX* allele variation within the clade.

Analysis of sequences orthologous to *Ra_g19519* from members of the Subgenus *Cylactis* (*R. arcticus*, *R. lasiococcus*, and *R. pubescens*) identified non-synonymous amino acid substitutions within conserved regions of *WOX*-like proteins. These substitutions may underlie the smooth phenotype observed in *Cylactis* species (Supplemental Figure 3).

While investigating *WOX* genes in *Rubus* we noted a few unexpected gene absences (Figure 3F). In the *R. argutus cv.* Hillquist genome we were unable to find the *WUS* ortholog, but it was found in the *R. ulmifolius* cv. Burbank assembly. The surrounding genes are present on Chromosome 1 in the cv. Hillquist assembly indicating the absence is from a loss of the gene in *R. argutus* or an issue with the assembly. Similarly, we were unable to find the *WOX2* gene for *R. idaeus* in the cv. Anitra assembly but did find it in the cv. Joan J assembly. This in-depth analysis across multiple *Rubus* genomes and allelic variations provides significant insights into the molecular mechanisms underlying prickleless phenotypes and offers pathways for targeted breeding in commercial *Rubus* fruit crops.

### Transformation and confirmation of *S* gene

We identified specific alleles of the *RuWOX1*/*RiWOX1* gene that contribute to the prickleless trait observed in several modern blackberry and raspberry cultivars. Although prickleless raspberry and blackberry cultivars are available, many elite cultivars, including PW-D0760, remain prickled. To verify that *RuWOX1* is causative of the prickleless phenotype and demonstrate the potential to enhance commercial lines, we developed a gene-editing approach to knock out *RuWOX1* in PW-D0760. Since PW-D0760 is triplex for the *Ruwox1-1* allele (Figure 3E), editing just the single wild-type copy of *RuWOX1* could yield the desired prickleless trait.

We implemented two gene-editing strategies targeting the *RuWOX1* gene in PW-D0760. In the first approach, we designed three guide RNA spacers (PWsp3329, PWsp3330, PWsp3331) to target a conserved region of *RuWOX1*, ensuring all haplotypes in PW-D0760 would be edited (Figure 4A). For the second approach, we designed a specific spacer (PWsp3332) targeting only the wild-type copy of *RuWOX1* in PW-D0760, allowing precise knockout of only the functional allele (Figure 4A).

**Figure 4.**
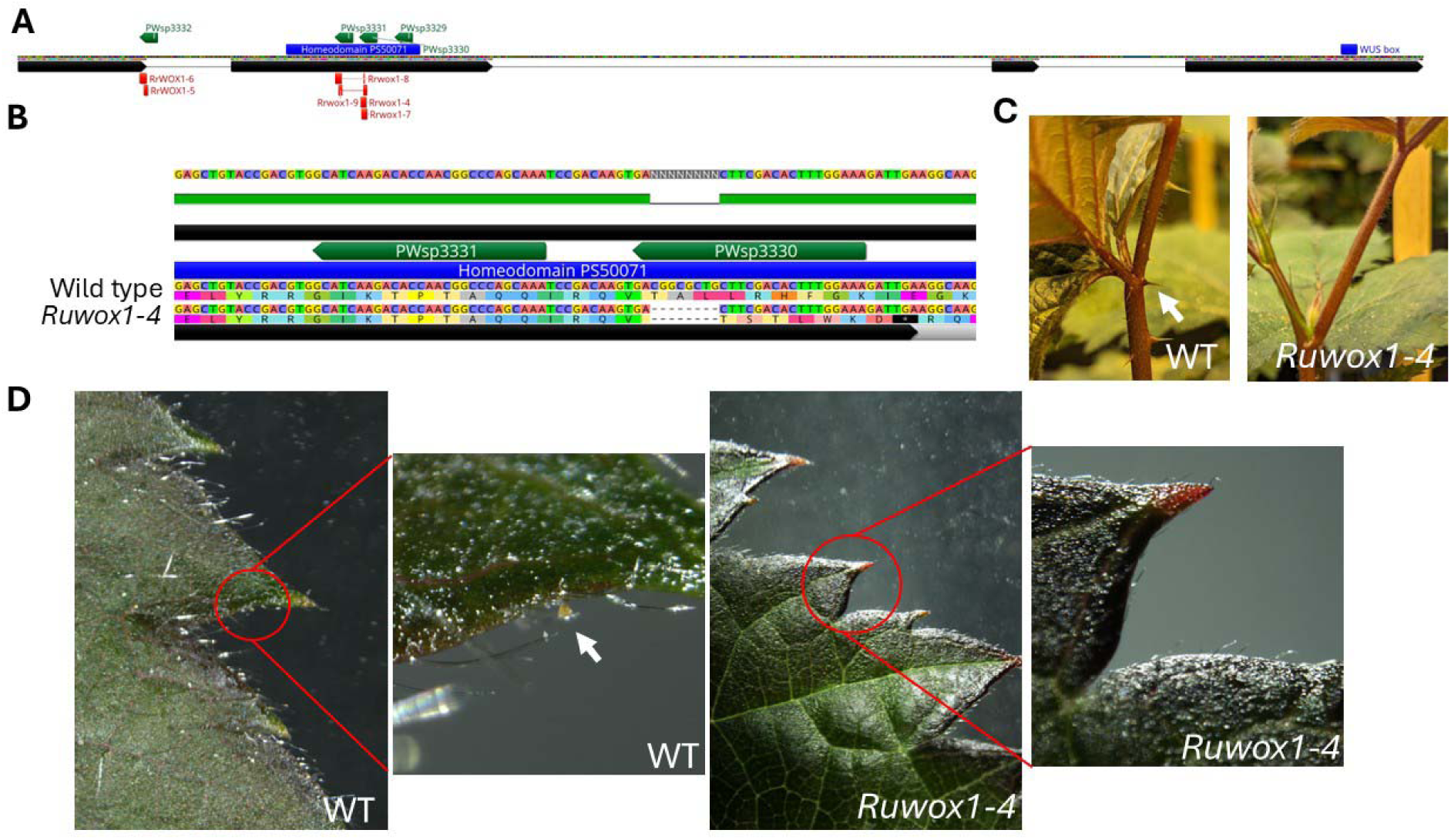
Edited alleles and mutant phenotype (A) *Rubus* wild-type *WOX1* gene structure. The spacer design is labelled in Green, the location of the mutant alleles are labelled in Red. The location encoding for two domains, the Homeodomain and WUS box, characteristic of the *WOX* family is labelled in blue. (B) *Ruwox1-4* allele is aligned to wild-type sequence showing an 8bp deletion leading to a premature stop codon. (C) Absence of prickles with *Ruwox1-4* allele in PW-D0760. Arrow indicates prickle in wild-type PW-D0760. (D) Absence of glandular trichomes with *Ruwox1-4* allele in PW-D0760. Arrow indicates prickle in wild-type PW-D0760.

Since blackberry had not previously been transformed, we also developed a transformation system specifically for this purpose. Through these strategies, we successfully created several edited alleles in PW-D0760. The first *Rubus* subg*. Rubus* edited allele, named *Rrwox1-4*, involved an 8bp deletion within the homeodomain, resulting in a premature stop codon within this critical region (Figure 4B). The *Rrwox1-4* plants exhibited both prickleless and glandular trichome-free phenotypes, confirming the targeted gene’s influence on these traits (Figure 4C, D).

In addition to *Rrwox1-4*, we generated five additional alleles in PW-D0760, each carrying mutations that introduced premature stop codons within or before the homeodomain motif (Table 6; Figure 4A). All edited plants exhibited the prickleless, glandular trichome-free phenotype, without any other notable phenotypic differences compared to the prickled, wild-type PW-D0760. Both edited and wild-type PW-D0760 plants maintained simple trichomes, suggesting that *RrWOX1* selectively influences the development of prickles and glandular trichomes without impacting other trichome types.

**Table 6:** **Alleles in Blackberry *WOX1***

| NAME | SOURCE | POLYMORPHISM | LOCATION WITHIN GENE |
| --- | --- | --- | --- |
| **Ruwox1-1<sup>a</sup>** | *R. ulmifolius* cv. Merton thornless | 8bp insertion | Homeodomain encoding region |
| **Ruwox1-2<sup>b</sup>** | *R. ulmifolius* cv. Burbank thornless | 4bp insertion | WUS box encoding region |
| **Riwox1-3<sup>c</sup>** | *R. idaeus* cv. Burnetholm | 8bp insertion | 2nd exon |
| **Rrwox1-4<sup>d</sup>** | *Rubus* subg. *Rubus* | 8bp deletion | Homeodomain encoding region |
| **Rrwox1-5<sup>D</sup>** | *Rubus* subg. *Rubus* | 6bp deletion | 1st exon/intron boundary |
| **Rrwox1-6<sup>D</sup>** | *Rubus* subg. *Rubus* | 10bp deletion | 1st exon |
| **Rrwox1-7<sup>D</sup>** | *Rubus* subg. *Rubus* | 8bp deletion | Homeodomain encoding region |
| **Rrwox1-8<sup>D</sup>** | *Rubus* subg. *Rubus* | 9bp/1bp deletions | Homeodomain encoding region |
| **Rrwox1-9<sup>D</sup>** | *Rubus* subg. *Rubus* | 5bp/5bp deletions | Homeodomain encoding region |

<sup>a</sup> s allele in modern prickless blackberries
<sup>b</sup> s allele retained in the unphased Burbank thornless genome assembly
<sup>c</sup> s allele in modern prickless red raspberries
<sup>d</sup> Novel s alleles created through gene editing resulting in the prickless phenotype in the tetraploid blackberry PW-D0760

### Alternate targets in blackberry

Previous GWAS and RNA expression analyses of a segregating population for the *s^ri^* allele in red raspberry identified several candidate genes for regulating the prickle trait (Khadgi and Weber, 2020b; Khadgi and Weber, 2020c). Among these, a primary candidate was MYB16-like, a transcription factor belonging to the MIXTA-like R2R3-MYB family (Ra_g19351), known to regulate the outgrowth of conical cells and the initiation of trichomes (small hair-like structures) in various plant species. However, in our study, sequencing of prickleless blackberry and red raspberry genomes revealed no loss-of-function mutations in MYB16-like coding regions.

Furthermore, gene knockout experiments showed no observable phenotypic differences in homozygous plants for this gene (data not shown), leading us to exclude *MYB16-like* from further analysis. Previous work characterizing genetic control of prickle differentiation in Rosaceae has provided some support for an overlap between regulation of prickle development and trichome development. In *Rosa hybrida*, orthologs of core transcription factors regulating trichome development in Arabidopsis were found to have higher expression in prickles than in bark tissue. Expression of several of these transcription factors, including *GL1*, *GL3*, *TTG1*, and *TTG2* were determined to be higher in bark of rose with a prickled phenotype than in bark of prickle-free rose(Swarnkar et al., 2021). To test whether these key regulators of trichome differentiation play an essential role in *Rubus* prickle formation, we identified orthologs of *AtGL1* (Ra_g21218) and *AtTTG2* (Ra_g29612) in blackberry (Supplemental Figure 4A, B). In both cases, we identified a single ortholog in PW-D0760. We designed guide RNAs to target these genes and used the previously described transformation method to recover edited blackberry events. We recovered events from the testing strategy for both genes which showed four unique loss-of-function alleles, indicating complete knockout of *RuGL1* and *RuTTG2*. These *Rugl1* and *Ruttg2* loss-of-function mutants both retained prickles, glandular trichomes, and simple trichomes, on leaves and canes. These results indicate that loss of these transcription factors required for trichome development in Arabidopsis is not sufficient to prevent development of any of these epidermal outgrowths in *Rubus*. In contrast, we also transformed PW-D0760 with a construct containing a constitutively repressive *p35S::GL1-EAR* fusion, and we recovered T0 events which lacked prickles and glandular trichomes but retained simple trichomes (Supplemental Figure 4C). This result suggests that the core trichome regulating factors in Arabidopsis have a role in prickle differentiation in *Rubus*, but functional orthology of members of these transcription factor families has diverged between Arabidopsis and *Rubus*.

## Discussion

We have identified the genetic basis underlying the prickleless trait in both red raspberry and blackberry. Previous efforts to map the genetic determinants of prickles in *Rubus* using single reference genomes resolved the locus to Chromosome 4 (Khadgi and Weber, 2020b; Johns et al., 2025). We refined and found the locus by integrating multiple genomes and a combination of genome-wide association study (GWAS), bulked segregant analysis (BSA), and inheritance-by-descent (IBD) analyses.

We created a new genome assembly of one of the first prickleless blackberries, *R. ulmifolius* cv. Burbank Thornless, and used several other *Rubus* genomes including, blackberry (*Rubus argutus*, cv. Hillquist (Brůna et al., 2022)), red raspberry (*Rubus idaeus*, cv. Anitra (Davik et al., 2022)) and black raspberry (*Rubus occidentalis* ‘ORUS 4115-3’ (VanBuren et al., 2016)) in our analyses. We used *Rubus argutus*, cv. Hillquist as the reference genome to find candidate mutations in the *S* locus (Brůna et al., 2022). By employing a diverse array of genetic mapping techniques, including GWAS, BSA and IBD analysis, we refined the location of the prickleless trait to a specific region on chromosome 4. Previous work had performed a GWAS on a red raspberry cross (prickle-free Joan J (*ss*) and prickled Caroline (*Ss*)) had located the *S* locus to a 1604kb physical region (Khadgi and Weber, 2020b). We took advantage of our diversity panel to combine information from both red raspberry and blackberry accessions. This depended on the knowledge that the *S* locus is allelic in blackberry and red raspberry (Lewis, 1939; Jennings, 1986; Jennings, 1988). The advantage of BSA and IBD analyses is the ability to use SNPs to identify precise recombination boundaries to delineate the region. Using the combination of genomic analyses on both sources of the *S* allele reduced the size of the physical region to less than 400kb.

We used RNA sequencing (RNA-seq) of blackberry plants segregating for the prickleless and prickled phenotype to prioritize potential candidate genes within the 400kbp region. Expression analysis has been successfully used to identify causative genes for mutants (Sommer et al., 1990). One disadvantage is that most mutant alleles are not null for RNA expression. We generated a self-crossed PW-D0007 population segregating for prickled and prickleless progeny. Of the 62 genes within our identified region, five genes had altered expression.

One of the five differentially expressed genes within the *S* locus was Ra_g19519, which was annotated as belonging to the *WUSCHEL-Related Homeobox* family. There was a 2-3 fold reduction in expression of Ra_g19519 in the RNAseq experiment, which could be explained by nonsense-mediated decay of the mutated allele (Hug et al., 2016) . The *WUSCHEL-Related Homeobox* (*WOX*) family constitutes a phylogenetically conserved group of homeodomain-containing transcription factors pivotal to plant developmental processes and cellular differentiation (Graaff et al., 2009). Named after the archetypal *WUSCHEL* (*WUS*) gene, which is essential for stem cell niche maintenance within the shoot apical meristem, *WOX* genes regulate meristematic activity, organogenesis, and spatial patterning across various plant tissues (Laux et al., 1996). WOX proteins possess a highly conserved homeodomain, facilitating DNA-binding capabilities that enable precise transcriptional control over downstream target genes critical for meristem functionality and organ development. We found that Ra_g19519 was most closely related to the Arabidopsis gene *AtWOX1*, AT3G18010, within the *WOX* family.

Using sequence similarity and synteny we showed the most direct ortholog of *AtWOX1* in blackberry is Ra_g20143. Within the Rosaceae, the subclade of Ra_g19519 is present in *Fragaria*, *Rosa* and *Rubus* but not in *Prunus*. This suggests an expansion of the *WOX1* subclade within the Rosoideae subfamily of the Rosaceae. In *Rosa hybrida,* gene expression has been studied in epidermal cells at different stages of prickle development (Swarnkar et al., 2021). The ortholog of a candidate gene in the blackberry locus, *RhWOX1(RcHm_v2.0_Chr7g0178511),* was ∼20x more highly expressed in the epidermis of growing prickles samples compared to initiating epidermis or hard prickle (Supplemental Figure 5). *AtWOX1* acts as a transcriptional repressor promoting cell proliferation during leaf initiation and leaf lamina growth(Ikeda et al., 2009)and is considered to act via the auxin biosynthesis pathway (Zhang et al., 2011; Nakata et al., 2018; Zhang et al., 2020b). Interestingly, *AtWOX1* is genetically redundant to *AtWOX3* and recent mapping in eggplant (*Solanum melongena*) has suggested a mutation in an *AtWOX3* ortholog is responsible for loss of prickles in the 17C02 cultivar (Nakata et al., 2012; Qian et al., 2021). In eggplant there has been a duplication of the *AtWOX3* related subclade although the roles of the expanded *SmWOX3* genes is unclear. In rice (*O. sativa*), the functional gene conferring the formation of trichomes was reported to also encode an AtWOX3 related protein (Sun et al., 2017). In *Zea mays*, the *NARROW SHEATH* (*NS1/NS2*) WOX genes define lateral domains of the leaf primordium, specifying marginal founder cells that contribute to saw-tooth hair and margin development (Nardmann et al., 2004) . In cucumber, *CsWOX3* negatively regulates fruit spine formation by repressing cell proliferation at the spine base (Xu et al., 2024) . In citrus, *CsWUS*, a *WOX* family member, modulates axillary meristem fate, controlling the developmental balance between thorn and branch formation (Zhang et al., 2020a) . Together, these examples illustrate how *WOX* genes are deployed to define specific lateral, epidermal, or meristematic domains in plant organs.

Manual annotation within the *S* locus led to the identification of mutations within Ra_g19519 orthologs RuBurbank_v1_000000F_038, in blackberry and Ri04_g21623, in red raspberry. Our findings reveal that an 8bp insertion in the second exon of *RuWOX1*, termed *Ruwox1-1*, introduces a premature stop codon in the homeodomain, likely leading to a non-functional protein and the subsequent prickleless phenotype. We also found an 8bp insertion in red raspberry, termed *Riwox1-2*, just after the homeodomain in Ri04_g21623 that also likely leads to a non-functional protein. The *Ruwox1-1* and *Riwox1-2* alleles co-segregate with the prickleless (*s^ru^*, *s^ri^*) alleles across the respective populations analyzed, strongly suggesting that these mutations are responsible for the loss of prickles.

The *R. ulmifolius* cv. Burbank Thornless genome provided additional insights into the genetic complexity of the prickleless trait. In an earlier GWAS of 374 blackberry genotypes grown by the University of Arkansas System Division of Agriculture, Burbank Thornless was the only prickleless genotype that was heterozygous for the SNP most strongly associated with prickleless canes at 33,636,565 bp on chromosome Ra04 (Johns et al., 2025) . We found that Burbank Thornless carries two distinct *RuWOX1* alleles: *Ruwox1-1* and *Ruwox1-2*, the latter characterized by a 4bp deletion within the WUS box and other SNPs. This compound heterozygosity may explain the thornless phenotype in this cultivar, despite heterozygosity at the *RuWOX1* locus, which typically results in a prickled phenotype. These findings highlight the importance of understanding the interaction between different alleles at the *RuWOX1* locus, as they have significant implications for *Rubus* breeding programs aiming to develop prickleless cultivars. Because the Burbank Thornless assembly is unphased, only one haplotype at the *S* locus was represented in the final reference sequence. In this case, the assembled haplotype corresponds to *Ruwox1-2*, while the alternative allele (*Ruwox1-1*) was carried forward into modern prickleless breeding germplasm. This underscores how reference assemblies may not always capture the functionally relevant allele and highlights the importance of haplotype-resolved assemblies for connecting reference genomes to functional alleles.

In the commercial blackberry line PW-D0760, the presence of the *Ruwox1-1* allele in triplex provided further validation of the gene’s role in prickle formation. Our gene-editing experiments in PW-D0760, designed to knock out *RuWOX1*, were successful. We generated several alleles carrying premature stop codons, with all resulting plants exhibiting the prickleless and glandular trichome-free phenotypes. These edited lines lacked glandular trichomes, suggesting that *RuWOX1* may regulate both prickle and glandular trichome development. However, no other phenotypic differences were observed between wild-type and edited plants, indicating that *RuWOX1* has a specific role in prickle and glandular trichome formation without affecting other trichome types or overall plant morphology.

Within our diversity panel, varieties that exhibited prickles always had glandular trichomes suggesting that these epidermal outgrowths share common developmental pathways. Our work indicates that *RuWOX1* could be a key regulatory gene influencing both traits. However, varieties that had glandular trichomes had a moderate correlation with the presence of prickles (r = 0.47). This suggests other factors are involved with prickle formation and further research is necessary to fully understand the genetic networks governing the formation of prickles and glandular trichome development.

Our findings provide a clear case for parallel evolution at the molecular level, where the prickleless trait arose from at least three different natural mutations (*Ruwox1-1*, *Ruwox1-2*, *Riwox1-3*) all in the same *WOX1* gene, even in distantly related *Rubus* species. This repeated independent targeting suggests *WOX1* acts as a key developmental hotspot for this trait. Such parallel evolution is often favored when a single gene acts as a developmental chokepoint with low pleiotropy, where it represents an efficient evolutionary pathway to alter a specific trait without incurring major fitness penalties. Our own gene-editing results support this, as the *Rrwox1-4* knockout plants were phenotypically normal, aside from the targeted loss of prickles and glandular trichomes. The reduced risk of pleiotropy makes *WOX1* an excellent target for crop improvement. At the same time. A recent study in *Solanum* revealed that prickle loss has repeatedly evolved through mutations in a *LONELY GUY* (*LOG*) cytokinin biosynthesis gene, known as *prickleless* (*PL)* (Satterlee et al., 2024) . At least 16 independent *log* mutations were identified across wild and domesticated species, and similar mechanisms were found in unrelated plants like barley, rice, jujube, and rose. The *PL* gene was shown to be a conserved ortholog shared across angiosperms, including *Rubus*.

In Arabidopsis a member of the *LOG* gene family, *AtLOG4*, gene has been shown to promote expression of *AtWUS* in the shoot apical meristem (Chickarmane et al., 2012) . A similar genetic interaction between *RuLOG* and *RuWOX1* may be conserved in glandular trichomes and prickles meristem formation. Gene editing of *PL* successfully removed prickles from eggplant without affecting other traits, highlighting a convergent, genetically simple mechanism that could be applied in *Rubus* breeding (Satterlee et al., 2024) . While *WOX1* is a confirmed pathway in *Rubus*, other candidate gene targets such as *LOG* could be examined to achieve loss of prickles in *Rubus*, possibly without loss of glandular trichomes.

This research provides significant insights into the molecular mechanisms underlying the prickleless trait in *Rubus* and demonstrates the potential of genome editing technologies in *Rubus* breeding. By precisely targeting the *RuWOX1* gene, we successfully created prickleless plants without the need for traditional breeding techniques, which often result in the loss of other desirable traits. Our study offers a promising pathway for the development of prickle-free commercial cultivars, which would greatly improve harvesting efficiency and reduce labor costs in the *Rubus* industry. Genome editing may be especially advantageous for the development of prickleless blackberries because of its recessive inheritance and the disadvantageous traits in linkage with the *s^ru^* locus, including susceptibility to winter injury, trailing growth habit, late harvest season, large seed size, and high fruit acidity (Moore, 1984). An extensive linkage disequilibrium block was discovered around the *s^ru^* locus in the University of Arkansas System Division of Agriculture blackberry germplasm, which indicates a precipitous loss of diversity in this region during decades of intensive selection for prickleless plants and suggests that linkage drag in the region may be difficult to overcome. Using gene editing, breeders can modify the prickled allele(s) at the *S* locus in an elite prickled variety with exceptional fruit quality and agronomic traits without the time and expense of further crossing and the complications of potential inbreeding and linkage drag in the distal portion of chromosome 4.

Future work should focus on further characterizing the regulatory role of *RuWOX1* in epidermal development and determining how this gene interacts with other pathways involved in trichome and prickle formation. Additionally, the development of more robust transformation systems for *Rubus* species will facilitate the broader application of gene-editing technologies, accelerating the development of improved cultivars with desirable traits such as pricklelessness, disease resistance, and enhanced fruit quality.

## Material and Methods

### Vector design

The binary plasmid, pWISE8807, was designed to express the *Lachnospiraceae bacterium* ND2006 Cas12a (LbCas12a) endonuclease (Zetsche et al., 2015) and 3 guide RNA spacers (PWsp3330: CCTCGCGCGGTGGTTCTGAAACC, PWsp3331: CTGGGCCGTTGGTGTCTTGATGC, PWsp3329: CCTCGCGCGGTGGTTCTGAAACC) to knock out the functional copy of the *WOX* gene in *PW-D0760* . Spacers were designed to mutate Exon I for the following reasons: (1) It is the first exon so frameshifts early in the gene are likely to produce loss of function phenotype; (2) This is the same region of the gene that the non-functional mutations in the other copies of *WOX* can be found. Sequence alignments of the exon between the four copies of the gene were completed to identify the optimal region to target for loss of function. Guides were expressed from the 3’ UTR of the *Ds.RED-Express2* gene (Strack et al., 2008) driven by the *OsAct1* promoter. Transcription of guide RNAs was terminated with the *StPinII* terminator. The guide cassettes were synthesized by GenScript and inserted in the binary plasmid with the *LbCas12a* expression cassette. A disarmed *Agrobacterium tumefaciens* strain was used to introduce a T-DNA from the binary plasmid. The T-DNA cassette expresses a plastid-targeted *NptII* expression cassette as a plant selectable marker, the cassette for LbCas12a endonuclease, *DsRED-Express2* with 3’UTR guide RNA cassette and a *ZsGreen* expression cassette for transgenic color marker selection. The binary plasmid was re-extracted from *Agrobacterium* and verified by whole plasmid sequencing by plexWell PRO™ (seqWell, Beverly, MA, USA).

### Plant Transformation

Blackberry nodal explants were harvested from the greenhouse and sterilized using ethanol and bleach, followed by sterile water rinses. Sterile nodes were then cultured on a propagation medium for 3 to 6 weeks prior to transformation. Explants excised from in vitro shoot cultures were inoculated with *Agrobacterium* prior to co-culture on sterile filter paper for 5-6 days and were then transferred to a selective medium. Two transformation methodologies were developed in which initial regenerants were usually chimeric, containing a mixture of non-transformed and transformed edited cells. To obtain uniformly-edited plants, an additional round of regeneration was induced *in vitro* from various chimeric tissues resulting in the recovery of edited plants suitable for characterization. We refer to this process as “chimera cleanup.” Using this approach, multiple uniform and uniquely-edited plants could be recovered from each chimera. Up to six uniquely-edited plants have been recovered from the cleanup of a single chimera. Time in culture post-transformation ranged between 3.5 to 8 months before plants suitable for characterization were rooted and acclimated to greenhouse conditions. The detailed information about transformation are in PCT/US2025/032956 and PCT/US2025/033708.

### Trichome and Prickle Phenotyping

Accessions used in the study were accessed through the USDA-ARS National Clonal Germplasm Repository (NCGR) at Corvallis, Oregon (Supplemental table 1). Leaves with petioles attached were collected from all included accessions and observed under a dissecting microscope. Accessions were grouped into four classes based on presence and abundance of various trichome types: (A) simple trichomes only, (B) simple trichomes and short (<1mm in length) glandular trichomes, (C) simple trichomes, short glandular trichomes and sparse elongated (>1mm in length) glandular trichomes, or (D) simple trichomes, short glandular trichomes, and abundant (>20 / cm of tissue) elongated glandular trichomes.

Presence/absence of prickles on petioles was also assessed. We observed variation in epidermal structures, where many lines had thin, non-lignified epidermal outgrowths we termed “bristles.” Under micropscopy, we observed that these bristles were elongated glandular trichomes, which could be differentiated from prickles by their thickness and presence of a glandular head. We scored plants according to large epidermal outgrowth types, assigning plants into one of four groups: (1) lack of “bristles” or prickles, (2) “bristles” only, (3) prickles only, or (4) both “bristles” and prickles. Prickle observations on petioles were cross-checked against observations from canes, and observation of a prickle in either organ resulted in the line being grouped as prickled.

### *Rubus ulmifolius* cv. Burbank Thornless genome sequencing, assembly, and annotation

The diploid *R. ulmifolius* cultivar ‘Burbank Thornless’ (PI 554060) was obtained from the USDA-ARS NCGR for genome sequencing. Nuclear flow cytometry with *Vinca major* as a standard estimated the genome size at ∼341 Mb. High-molecular-weight DNA extracted from young leaves was sequenced with PacBio Sequel (six SMRT cells). Illumina Hi-C libraries were prepared from young leaf tissue, while RNA from root tips, leaves, and stems was used for Illumina RNA-seq and PacBio Iso-Seq. PacBio long reads were assembled with FALCON/FALCON-Unzip and under-collapsed heterozygosity was resolved with Purge Haplotigs (Chin et al., 2016; Roach et al., 2018). Scaffolding was performed with Hi-C data using Juicer/3D-DNA followed by manual polishing with Juicebox Assembly Tools (Durand et al., 2016; Dudchenko et al., 2017; Dudchenko et al., 2018). The final assembly comprised seven chromosome-scale scaffolds, which were ordered and oriented against the *R. argutus* cv. Hillquist reference genome.

For structural annotation, transcript assemblies were generated from RNA-seq and Iso-Seq data and combined with protein homology from 24 diverse plant species and curated databases. Gene models were predicted using multiple ab initio and evidence-based approaches (AUGUSTUS, FGENESH, EXONERATE, PASA) and refined by PASA to add UTRs, splice site corrections, and alternative isoforms(Salamov and Solovyev, 2000; Slater and Birney, 2005; Stanke et al., 2006). Low-confidence models overlapping repeats or lacking transcript/homology support were filtered. Functional annotation was assigned using BLAST searches against SwissProt, Araport11, and NCBI databases, InterProScan domain analysis, and KEGG/eggNOG orthology (Zdobnov and Apweiler, 2001; Moriya et al., 2007; Camacho et al., 2009; Huerta-Cepas et al., 2017; Afgan et al., 2018). Complete methods describing genome assembly and annotation are provided in Supplemental File 1.

### Genome-wide association study

The software package GWASpoly was used to perform genome-wide association mapping (Rosyara et al., 2016). Genomic DNA was extracted with Qiagen DNeasy Plant Mini Kit and sequencing were performed with the NovaSeq 6000 instrument with 150bp paired end reads to approximately 30X coverage. Initially, reads were cleaned with HTStream, mapped to the *Rubus argutus* cv. Hillquist reference genome with the aligner “bwa mem” (Li and Durbin, 2010), and variants were called with freebayes in tetraploid mode “-p 4” (Garrison and Marth, 2012). The variant set was randomly downsampled to 1 million SNPs then converted to allele dosage format with the GWASpoly function ‘VCF2dosage’ the following filters: min.DP=2, max.missing = 0.05, min.minor=5. After filtering, 333,198 SNPs remained for analysis. Marker significance scores were computed with the ‘GWASpoly’ function using the ‘1-dom’ tetraploid model with binary prickle/no-prickle trait scores and *p*-values were adjusted with a Bonferroni correction. The analysis included 29 prickleless and 14 prickle control samples. Given that the reference genome (Hillquist) contains the dominant allele for the prickle trait, the 1-dom-ref marker scores were used for the genome scan Manhattan plot.

#### BSA

We generated a segregating population by self-crossing PW-D0007, a line which was determined to be triplex for the ‘*s*’ allele (*Ssss*). The resulting progeny displayed either the prickleless (32 individuals) or prickled phenotype (65 individuals). We used this population in a bulked segregant analysis. For each bulk, DNA was extracted with DNeasy Plant Mini kit and pooled. High-throughput sequencing was then performed on both groups with the NovaSeq 6000 instrument with 150bp paired end reads. We generated ∼330X genome-wide sequence data representing the genetic makeup of the prickleless and prickled groups. The reads were mapped to the *Rubus argutus* cv. Hillquist genome assembly with the aligner ‘bwa mem’ (Li and Durbin, 2010) and variants were called with freebayes (Garrison and Marth, 2012). To identify genomic regions associated with the prickleless trait, the variants were filtered by allele frequency to match expectations of the recessive inheritance: prickleless bulk allele frequency >= 0.95 and control bulk allele frequency <= 0.85. The number of filtered variants per 100kbp window with a 50kb step size were then plotted to identify regions associated with pricklelessness.

#### IBD

Whole genome sequencing reads obtained for GWAS were aligned to the *Rubus argutus* cv. Hillquist, *Rubus occidentalis* v3, and *Rubus idaeus* cv. Anitra genomes and visualized in the jbrowse genome viewer. Syntenic regions of the three genomes were identified initially by selecting several genes within the *Rubus argutus* cv. Hillquist genome and running BLAST with the other genomes as the subject. Regions containing the resulting BLAST hits were then aligned with the regions from the other genomes. Most gene models from the region of one genome were seen in the others with high sequence conservation. SNPs correlating with differences in the prickle phenotype were checked in the other genomes. Due to SNP density in the region and the sequence diversity among the cultivars a region of 20Kbp was scanned in each direction any time a breakpoint in one cultivar was assumed to reduce the chance of calling a false breakpoint. SNPs associated directly with the prickleless phenotype were required to be homozygous in the prickleless cultivars and no more than heterozygous in the prickled cultivars. Because the *Rubus idaeus* cv. Anitra is prickleless, SNP calling was adjusted when comparing in that context. The reverse complement of the region from the *Rubus occidentalis* was used when aligning the regions.

### Orthology search for candidate gene

The genomic and protein coding sequence of Ra_g19519 were used to find orthologs in *Prunus avium* (Shirasawa et al., 2017), *Rubus occidentalis* v3 (VanBuren et al., 2016), *Rubus argutus* cv. Hillquist (Brůna et al., 2022), *Rubus ulmifolius* cv. Burbank thornless (this work), *Rubus L.* sub *Rubus* Watson (Paudel et al., 2025), *Rubus idaeus* cv. Anitra (Davik et al., 2022), *Rubus idaeus* cv. Joan J., (Zhou et al., 2023) and *Arabidopsis thaliana* Araport11 (Cheng et al., 2016). Additional searches using Arabidopsis *WOX* genes as query were conducted to find additional family members. Results from Hmmer3 (Eddy, 2008; Eddy, 2009; Eddy, 2011) and BLAST (Camacho et al., 2009) searches were aligned using MAFFT (Katoh et al., 2002; Katoh and Standley, 2016) and Clustalo (Sievers et al., 2011; Sievers and Higgins, 2018), for DNA and protein sequences, respectively. The DNA from conserved exons were selected including insertions and deletions to keep mutants close to their non-mutant relatives. UPGMA consensus trees were built with the alignments using a Jukes-Cantor distance model and 500 bootstrap resampling (Jukes and Cantor, 1969).

### RNA sequencing

For RNA analysis between our BSA prickles/prickleless samples, the tips of blackberry plants were excised in the greenhouse, immediately submerged in RNA Later (Invitrogen), placed on ice, and transported to a 4°C refrigerator, where they were stored for 24h to ensure thorough penetration of the RNA Later into the tissue. Meristems were then carefully dissected under a stereomicroscope and stored at −20°C until RNA extraction. RNA was extracted using the QIAGEN RNeasy Plant Mini Kit (QIAGEN) with buffer RLT, incorporating two apical meristems per sample. The extraction buffer was supplemented with 2.5% polyvinylpyrrolidone-40 (PVP-40) (PhytoTech Labs), and samples were incubated at 65°C for 10 minutes following grinding and buffer addition. An additional wash-step was included in the protocol, and RNA was eluted in 50 µL of nuclease-free water (Invitrogen).

HTStream (Petersen et al., 2015) was used to check reads for quality and prepare for mapping. Hisat2 (Kim et al., 2019) was used to map reads to the *Rubus argutus* cv. Hillquist genome. The lowest mapping rate for the samples was 84.5%. Htseq-count(Anders et al., 2015) was used to count transcript abundance as counts for prickleless and prickled samples were analyzed with DEseq2 (Love et al., 2014)

## Data availability

See supplemental file

## Supplemental Tables

Supplemental Table 1

Panel of 268 *Rubus* accessions, including 224 prickled and 44 prickleless accessions from eight subgenera, ensuring a broad representation of genetic and phenotypic variation for the prickle, trichome and glandular trichome traits. Names, plant introduction (PI) number, inventory number, taxon as listed in the USDA-ARS GRIN-GLOBAL database are listed.

Supplemental Table 2

Blackberry accessions in diversity panel with pedigree and predicted zygosity for *s* allele

Supplemental Table 3

Red raspberry accessions in diversity panel with pedigree and predicted zygosity for *s* allele

Supplemental Table 4

Predicted proteins in the *R. ulmifolius* cv. Burbank Thornless genome assembly along with physical positions, closest homolog among the *Rubus argutus* cv. Hillquist genes, and matches obtained from BLASTP analyses with NCBI nr, Araport11, RefSeq, SwissProt and TrEMBL databases as subjects.

Supplemental Table 5

Functional annotations of predicted transcripts in the *R. ulmifolius* cv. Burbank Thornless genome from InterProScan, Gene Ontology (GO), KEGG orthologs, and KEGG pathways.

Supplemental Table 6

Complete list of annotated 62-115 gene models found in mapped 324 Kb region Supplemental Table 7

Genes downregulated in global expression analysis between prickled and prickleless lines in the PW-D0007 blackberry self-crossed population. Expression analysis revealed 2357 up-regulated genes in the prickled samples and 1921 down-regulated in the prickled samples compared to the prickleless samples.

## Authors’ Contributions

B.A., T.P., A.S.C. and G.P. performed RNAseq and genomic analysis. C.O. coordinated and did phenotypic analysis of GWAS. L.R., and A.F. performed trait testing. J.R. and W.L. performed plant care. N.G. performed vector construction. E.D and D.C. performed transformation. S.L., P.M. and A.F. provided project governance. B.C.W.C. and G.P. authored the manuscript and provided conceptual guidance. R.A. performed DNA and RNA extractions, library preparation, and the PacBio genome assembly for Burbank thornless genome. O.D., M.P., D.W. and E.L.A. performed the *in situ* Hi-C experiment, Hi-C-guided assembly, and associated analyses. T.B. performed structural and functional annotation analyses. M.B. performed functional annotation analyses. A.S. assisted with data management and validation during annotation. N.B. coordinated sample collection and flow cytometry. D.M. coordinated and assisted with Hi-C sequencing, RNA-Seq, and genome assembly. M.W., D.J.S., D.M., E.A., and H.A. coordinated research and provided conceptual guidance. M.W. authored the supplemental file describing the Burbank Thornless genome assembly and annotation. All authors approved the final manuscript.

## Data Availability Statement

Raw PacBio and IsoSeq sequencing data and the genome assembly of *R. ulmifolius* presented here are available at the NCBI under Bioproject ID PRJNA1263752. Hi-C data are available on Bioproject PRJNA512907 (Biosample SAMN14122117; SRA SRX7735966). Illumina transcriptome data are available on Bioproject PRJEB36280 (BioSamples ERX5171502, ERX5171501, ERX5171495, ERX5171494). Interactive Hi-C contact maps of the Burbank Thornless genome sequence assembly are available via https://t.3dg.io/burbank-blackberry-Fig-1 and via the www.dnazoo.org website (https://www.dnazoo.org/assemblies/rubus_ulmifolius). The Burbank Thornless genome assembly and annotation can also be accessed at the Genome Database for Rosaceae (<u>Publication datasets | GDR</u>) under the accession number tfGDR1091 and on phytozome (https://phytozome-next.jgi.doe.gov/info/Rulmifolius_v1_1).

## Acknowledgments

We thank Kim Hummer and Jill Bushakra from the USDA-ARS, National Clonal Germplasm Repository, who assisted with sample collection and plant maintenance. We thank Gina Fernandez from NCSU who grew Burbank Thornless plants in her greenhouse and who participated in fruitful conversation and gave valuable advice at the inception of this project along with Felicidad Fernández-Fernández and Mario Caccamo from NIAB-EMR. We also thank Ryan Rapp, Aabid Shariff, and Xiaoyu Zhang from Pairwise for their assistance with IsoSeq analyses. We thank Mike Stratton and Julia Wilson for their continuing support for the 25 genomes for 25 years project. We thank Michelle Smith, Craig Corton, and Karen Oliver for processing RNA-seq samples. Genome assembly was performed in association with the DNA Zoo Consortium (www.dnazoo.org), which acknowledges support from Illumina, IBM, and Pawsey Supercomputing Center.

---

**Supplemental Figure 1.**
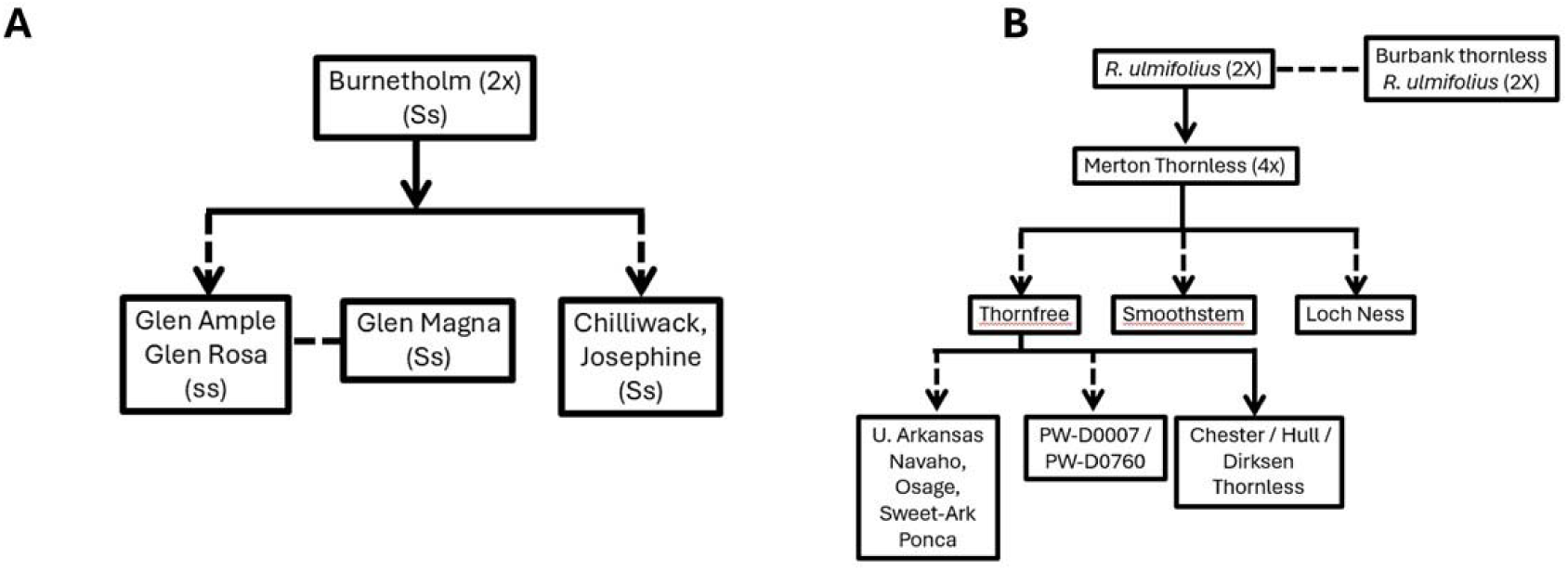
Pedigree of the *s* allele associated with the prickleless phenotype in commercial red raspberry (A) and blackberry (B) lines used for inheritance-by-descent models.

**Supplemental Figure 2.**
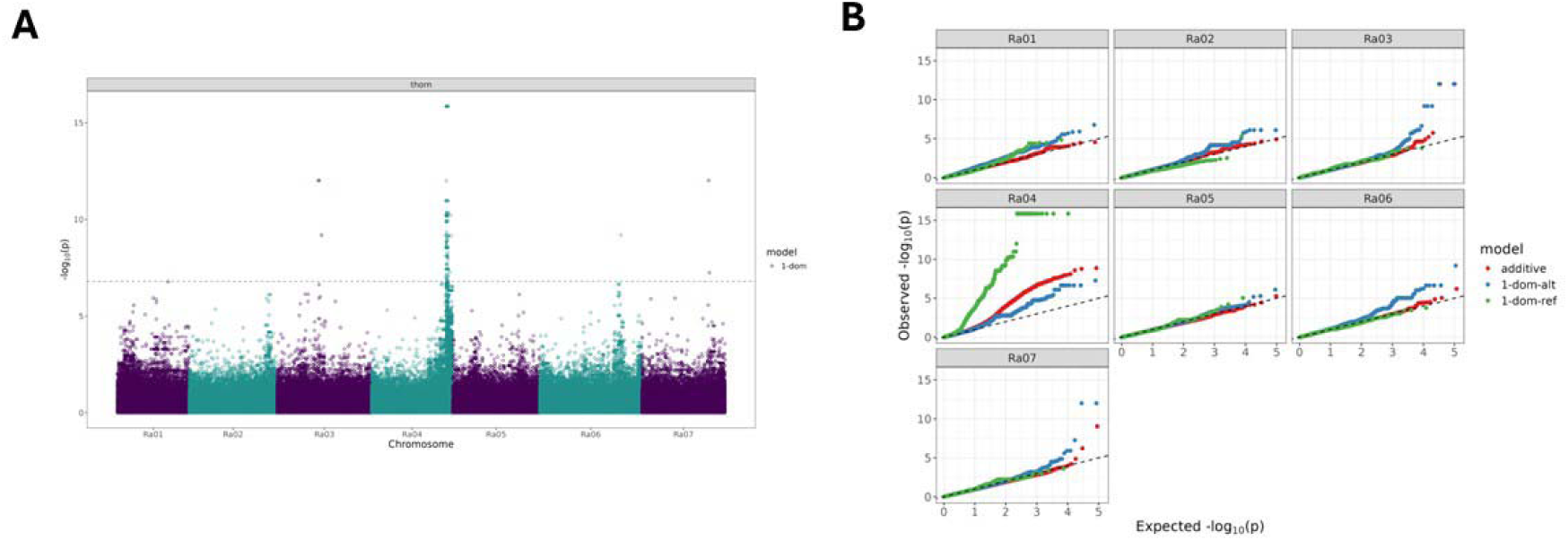
A. SNPpoly GWAS results for all chromosomes. B qqPlot

**Supplemental Figure 3.**
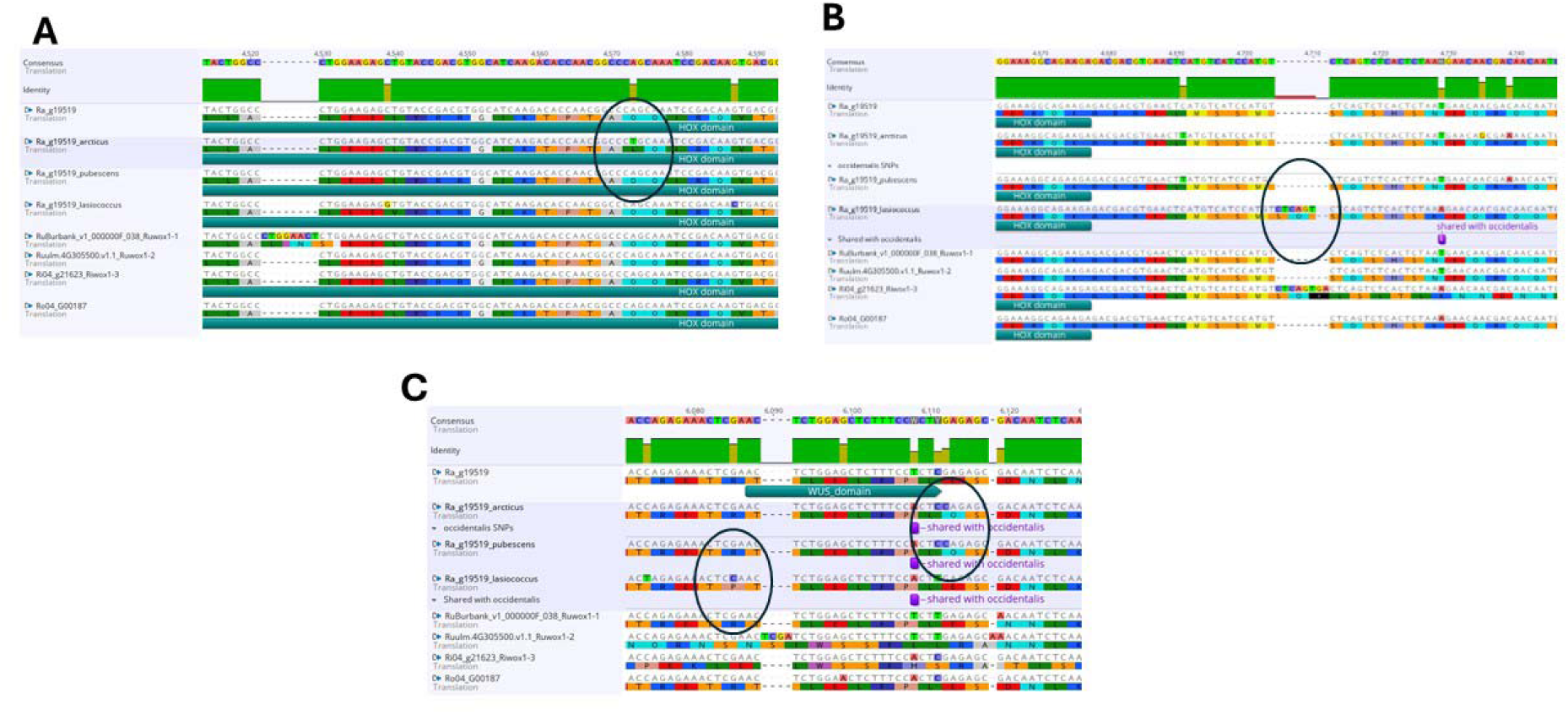
Analysis of sequences orthologous *to RaWOX1* (Ra_g19519) from *R. arcticus*, *R. lasiococcus*, and *R. pubescens* identified non-synonymous amino acid substitutions within conserved regions of WOX-like proteins. (A) Hydrophilic residue Q>L in smooth *R. arcticus* compared to other *Rubus* WOX1. (B) *R. Lasiococcus* has insertion near homeodomain in same position as seen in *Riwox1-3* but does not have a stop codon. (C) *R. arcticus, R. lasiococcus, and R. pubescens* all carry non-synonymous SNPs near the WUS box domain. These substitutions may underlie the smooth phenotype observed in these *Cylactis* species

**Supplemental Figure 4.**
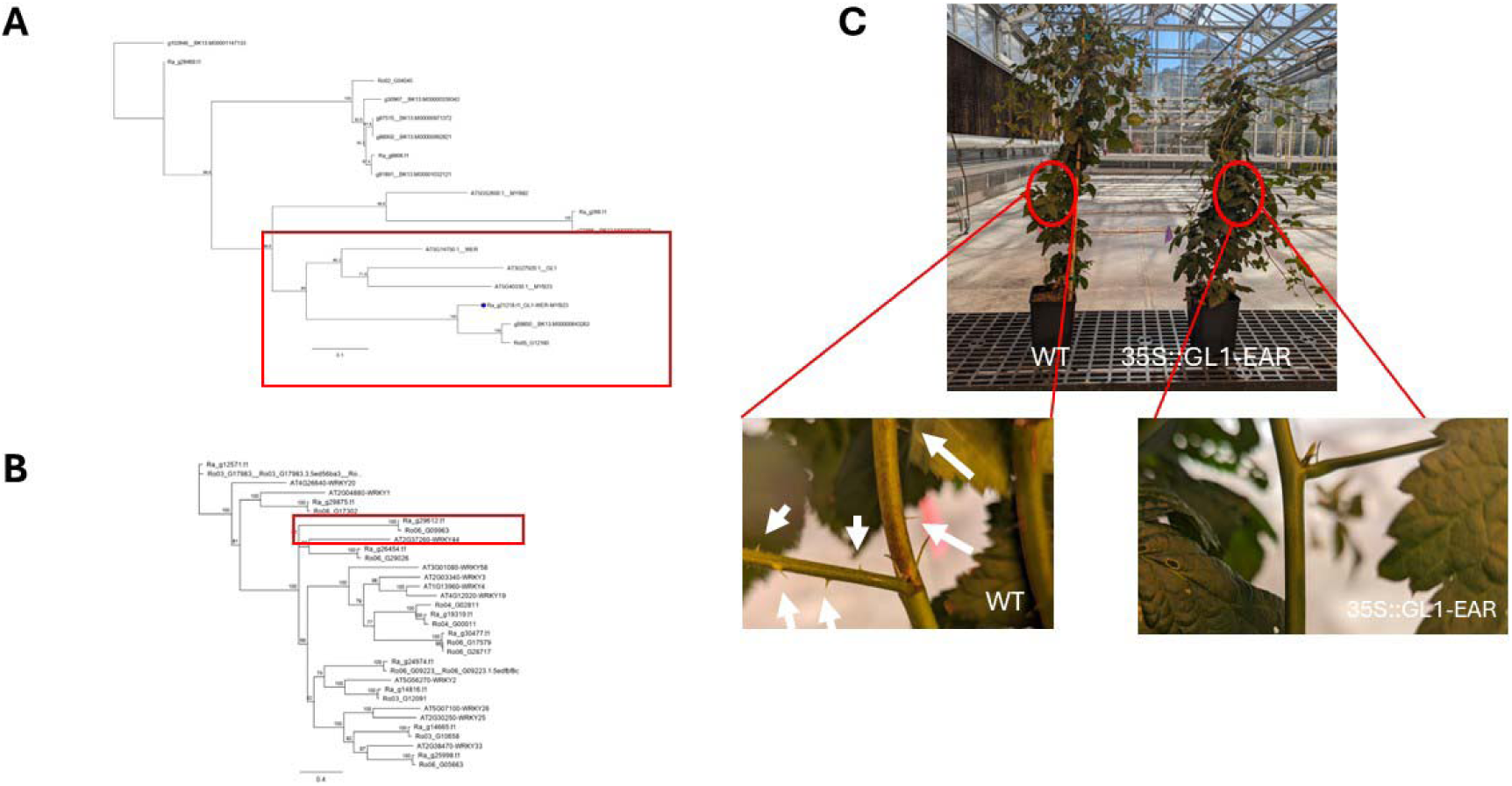
Alternate targets for prickleless in Blackberry. (A) Phylogenetic tree of orthologs in blackberry to the *AtGLABARA1* gene. The Red box highlights the *GL1* clade. (B) Phylogenetic tree of orthologs in blackberry to *the AtTRANSPARENT TESTA GLABROUS2* gene. The Red box highlights the *AtTTG2* clade. (C) Prickleless phenotype of PW-D0760 plants containing the *35S::GL1-EAR* fusion.

**Supplemental Figure 5.**
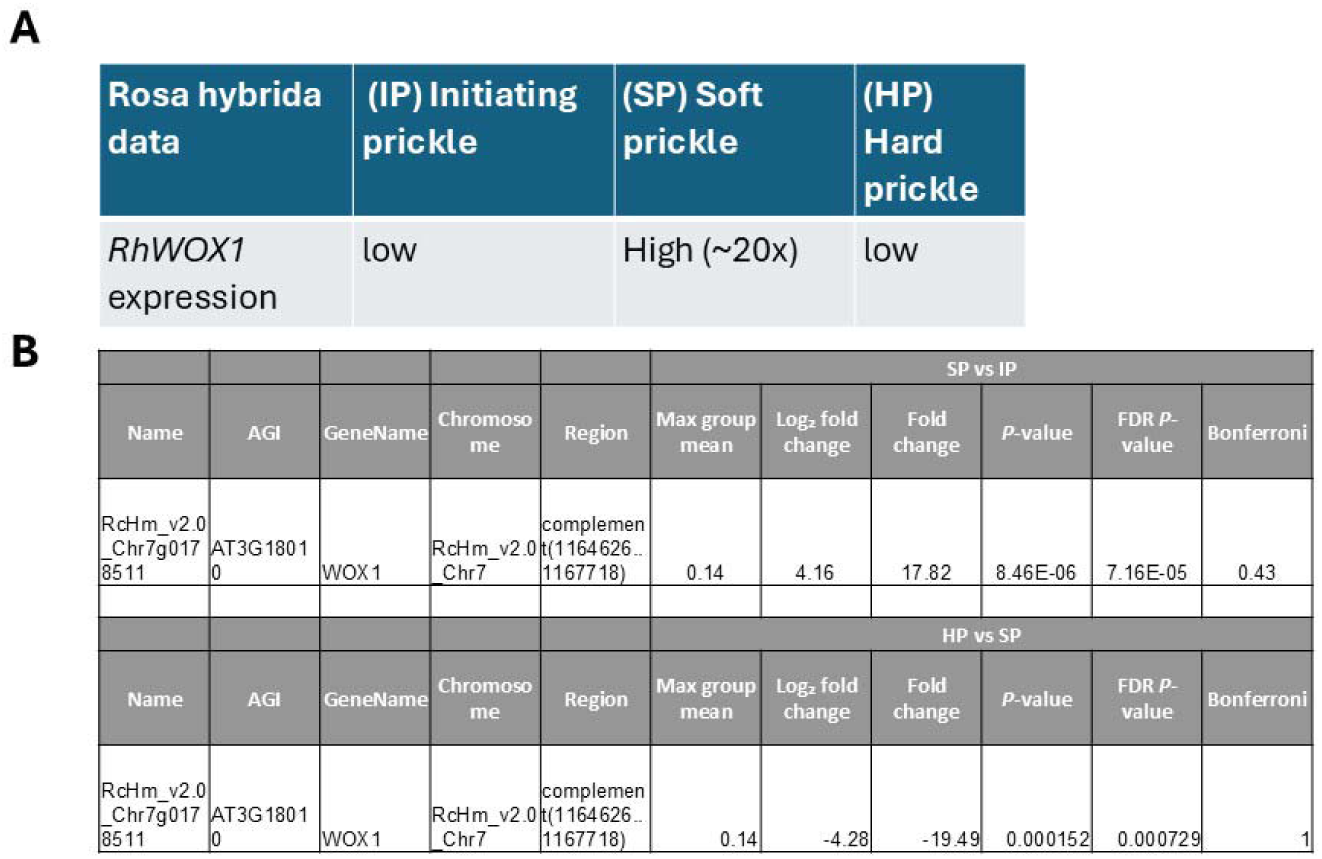
Expression data of *RhWOX1 (RcHm_v2.0_Chr7g0178511),* the *Rosa hybrida* orthologue of *RaWOX1,* in the epidermis cells at different stages of prickle development^29^. (A) *RhWOX1* was 20x more highly expressed in the epidermis of initiating prickles samples compared to initiating epidermis or hard prickle samples. (B) Data taken from Supplemental dataset 1 and 3 comparing differentially expressed genes from each tissue type. Initiating prickle together with bark (IP), soft prickle (SP), hard prickle (HP) - See more details in Figure 1. Swarnkar, M. K., Kumar, P., Dogra, V. & Kumar, S. Prickle morphogenesis in rose is coupled with secondary metabolite accumulation and governed by canonical MBW transcriptional complex. *Plant Direct* **5**, e00325 (2021).

