## Supplemental file 1 for "The control of prickle formation in *Rubus*"

**Supplemental File 1: Detailed methods and results for the *R. ulmifolius* cv. Burbank Thornless assembly and annotation**

**Materials and Methods**

***Plant material and genome size estimation***

The diploid *R. ulmifolius* cultivar Burbank Thornless (PI 554060) was sourced from the USDA National Clonal Germplasm Repository (NCGR) for genome sequencing and assembly. Leaf tissue was harvested from a single plant grown in the greenhouse at the USDA-NCGR, in Corvallis, Oregon for flow cytometry, DNA extraction, and PacBio and Hi-C sequencing. Burbank Thornless plants propagated by NCGR were sent to North Carolina State University (NCSU). Tissue from root tips and actively growing leaves and stems was harvested from greenhouse-grown plants at NCSU for RNA sequencing and IsoSeq. The DNA content and genome size of Burbank Thornless were estimated using nuclear flow cytometry with DAPI staining. Flow cytometry was performed using young, unexpanded Burbank Thornless leaves in biological triplicate with *Vinca major* as an internal standard.

***DNA extraction, library preparation, and sequencing***

High molecular weight DNA was extracted from young, unexpanded leaves of *R. ulmifolius* cv. Burbank Thornless following a modified CTAB method^1^ for PacBio sequencing. DNA quality was assessed with Pulsed Field Gel Electrophoresis (BioRad, Hercules, California), and quantified with a Qubit fluorometer (ThermoFisher Sci., Waltham, Massachusetts). Genomic DNA was sheared to achieve fragments in the 15–40 kb size range using a 26-gauge blunt end needle (ThermoFisher UK Ltd HCA-413-030Y GC Syringe Replacement Parts 26 g, 51 mm) and 1 mL luer-loc syringe. Sheared DNA was cleaned using 1X AMPure PB beads before library preparation. Fragments were enzymatically repaired and used to construct a long read (20 kb) PacBio Sequel genomic library with a SMRTbell™ Template Prep Kit 1.0-SPv3 following the manufacturer’s recommendations (Pacific Biosciences Inc., Menlo Park, CA, USA). The SMRTbell templates were size selected by BluePippin electrophoresis (Sage Science Inc., Beverly, MA, USA). Template DNA ranging from 15 to 50 kb in size was sequenced in six PacBio Sequel Single-Molecule Real-Time (SMRT) cells on a PacBio Sequel instrument at the NCSU Genomic Sciences Laboratory. Young leaf tissue from a Burbank Thornless plant subjected to 48 hours of darkness was collected for Hi-C sequencing. An *in situ* Hi-C library was prepared following an established protocol^2^ and sequenced on the Illumina NextSeq 500 as 80-bp paired-end reads.

***Genome sequence assembly***

A contig-scale assembly was generated with PacBio sequence data using FALCON and FALCON-Unzip software applications^3^. Error correction on the phased assembly was performed with the Arrow consensus model in the PacBio GenomicConsensus package using default parameters. The *k-mer* distribution of unassembled, corrected PacBio reads showed a bimodal distribution, indicating high heterozygosity. Therefore, under-collapsed heterozygosity was resolved using the Purge Haplotigs pipeline, by identifying syntenic pairs of contigs and moving one to a haplotig pool^4^. Hi-C data were aligned to the Purge Haplotigs draft assembly using Juicer v1.6.2^5^. The 3D *de novo* assembly (3D-DNA) pipeline^6^ was used to construct a candidate Hi-C-guided assembly and contact map. The assembly was then reviewed and polished using Juicebox Assembly Tools^7^. Interactive Juicebox.js [Robinson et al., 2018: PMID 29428417] contact maps before and after the Hi-C-guided assembly step are available for review at https://www.dnazoo.org/assemblies/rubus_ulmifolius. The 7 chromosome-length scaffolds were ordered and oriented to match the *R. argutus* blackberry assembly^8^ (Figure S1). Synteny of the Burbank Thornless genome to the *R. argutus* genome was assessed with LastZ^9^ with the following parameters: --notransition --step=20 --nogapped --format=maf --ambiguous=iupac --hspthresh=50000.

**Gene prediction and annotation**

*RNA extraction, library preparation, and sequencing*

Total RNA was extracted from three tissue types (root tips, actively growing leaves, and actively growing stems from same plant) using the Spectrum Plant Total RNA Kit (Millipore Sigma, Burlington, MA) following the manufacturer’s protocol. The purity and concentration of the RNA was assessed with a 2100 Bioanalyzer (Agilent Technologies, Santa Clara, CA). Samples determined to an RNA integrity number (RIN) value above 7.0 using a Qubit 4.0 fluorimeter (Thermo Fisher Scientific, Waltham, MA) were submitted for subsequent sequencing. Two duplicate RNA-Seq libraries were produced for leaf and stem tissue and sequenced with an Illumina HiSeq X instrument at Scientific Operations core at the Wellcome Sanger Institute. Total RNA from the leaf, stem, and root tissues were pooled and used for Iso-Seq library preparation. Standard PacBio Iso-Seq SMRTbell libraries were prepared by Genewiz (South Plainfield, NJ) and one SMRT cell was sequenced with Sequel II. Full-length transcripts were identified using the Iso-Seq 3 application in SMRTLink 5.0. Multiple reads of the same SMRTbell sequence or the subreads from the same polymerase read were combined to produce one high-quality circular consensus sequence (CCS). The CCS reads were then classified as full-length based on the presence of both cDNA primers and polyA tails. Full-length reads were further classified as chimeric or non-chimeric reads based on the presence or absence of primers in the middle of the sequences. Finally, the iterative clustering and error correction algorithm was used to extract and polish consensus isoforms to obtain high-quality and low-quality isoforms.

*Structural Gene Annotation*

Transcript assemblies were generated from ~30M pairs of 2 x 150 and ~65M pairs of 2 x 100 stranded paired-end Illumina RNA-seq reads. The reads were assembled using PERTRAN, which conducts genome-guided transcriptome short read assembly via GSNAP^10^ and builds splice alignment graphs after alignment validation, realignment, and correction. Approximately 1.3M PacBio Iso-Seq non-chimeric CCSs were corrected and collapsed by a genome-guided correction pipeline, which aligned CCS reads to the genome with GMAP^11^ with intron correction for small indels in splice junctions and clustered alignments based on intron structure or ≥ 95% overlap for single-exon transcripts. Approximately 69,000 full-length transcripts were obtained. Subsequently, transcript assemblies from both Illumina and PacBio data were integrated and refined using PASA^12^, yielding 87,104 transcript models prior to final filtering.

The genome was soft-masked with RepeatMasker^13^, using a species-specific repeat library. The repeat library consisted of de novo repeats predicted by RepeatModeler2^14^ generated on the *R. argutus* and *R. ulmifolius* genomes and common Viridiplantae and Embryophyta repeats in RepBase and Dfam. Putative gene loci were identified by transcript assembly alignments and protein homology evidence. Protein sequences from a diverse set of 24 plant species (Arabidopsis thaliana, Beta vulgaris, Cannabis sativa, Carya illinoinensis, Cucumis sativus, Fragaria vesca, Glycine max, Gossypium raimondii, Liriodendron tulipifera, Malus domestica, Medicago truncatula, Mimulus guttatus, Morus notabilis, Oryza sativa, Populus trichocarpa, Potentilla anserina, Prunus persica, Rosa chinensis, Solanum lycopersicum, Sorghum bicolor, Vitis vinifera, and Ziziphus jujuba) along with eukaryote proteins from Swiss-Prot (release 2022_04) were aligned to the repeat-soft-masked *R. ulmifolius* genome with EXONERATE^15^. Alignments were allowed to extend up to 2 kb beyond gene boundaries unless they overlapped with another locus on the same strand.

Gene models in each locus were predicted using multiple approaches, including FGENESH+ and FGENESH_EST^16^, EXONERATE, AUGUSTUS^17^, and homology constrained ORFs from the PASA transcript assembly. AUGUSTUS was trained on the high confidence PASA-derived ORFs and intron hints from short read alignments. For each locus, the highest scoring prediction was selected based on positive criteria, including EST and protein support, and one negative criterion: overlap with repeats. The selected gene predictions were improved by PASA, which added untranslated regions (UTRs) and alternative transcripts and corrected splice sites.

PASA-improved gene model proteins were evaluated for homology against the previously described proteomes to obtain Cscore and protein coverage metrics. The Cscore represents the ratio of BLASTP score for a given protein to the score of its mutual best hit, while protein coverage is the percentage of a protein aligned its best homolog. PASA-improved transcripts were selected based on a combination of Cscore, protein coverage, EST support, and coding sequence (CDS) overlap with repetitive elements. Transcripts with a Cscore and protein coverage of ≥ 0.5, or those supported by ESTs, were retained. For gene models with CDS regions overlapping repeats by more than 20%, more stringent criteria were applied: a minimum Cscore of 0.9 and homology coverage of at least 70%.

Selected gene models were further analyzed for protein domains using Pfam^18^. Gene models lacking strong transcriptomic and homology support and whose proteins had over 30% overlap with Pfam-annotated transposable element (TE) domains were removed. Additional manual curation was performed to exclude incomplete gene models, those with weak homology support and no full-length transcript evidence, short single-exon models (CDS <300 bp) lacking recognizable protein domains or expression support, and repetitive models unsupported by strong homology evidence. Gene-level macrosynteny with *R. argutus* was evaluated using GENESPACE, which identifies conserved gene order across genomes.

*Functional gene annotation*

Functional annotation of predicted gene models was performed by querying SwissProt, Araport11, NCBI nr, RefSeq and TrEmbl protein databases with BLAST+ blastp-fast algorithm^19^. The predicted protein-coding sequences of the 38,105 transcripts identified in the structural annotation were used as queries with an expectation value cutoff of 1e-3. BLAST+ searches were conducted on the Galaxy platform^20^ with locally installed databases except for Araport11, which was obtained from The Arabidopsis Information Resource (TAIR, <https://www.arabidopsis.org/>). Moreover, the closest homolog among the *Rubus argutus* Hillquist Genome v1.0 genes^8^ for each protein was determined with the same method. InterProScan v5^21^ was used to assign InterPro domains, and Gene Ontology (GO) terms to the predicted proteins. KEGG ortholog and pathway associations were inferred with KAAS (KEGG Automatic Annotation Server) v2.1^22^ using bi-directional best hit and eggNOG-mapper v2^23^, respectively.

**Results and Discussion**

***Chromosome-length genome assembly***

A combined total of 3.3 million PacBio post-filtered reads with an average length of 9,803 bp were generated from the six SMRT cells, resulting in a total of 32.5 Gb of sequence (~89X Genome Coverage) (Table 1). These reads were used to generate an initial FALCON-Unzip assembly comprised 426 Mb of sequence in 1131 contigs with an N50 of 915 kb and a maximum contig length of 9.1 Mb. After Purge Haplotigs was used to resolve under-collapsed heterozygosity, the optimized assembly consisted of 341 Mb assigned to 463 primary contigs with a contig N50 of 1197 kb and a maximum contig length of 9.1 Mb. The Hi-C library was sequenced to produce 107,483,588 million paired-end reads, totaling 17.2 Gb of sequence data. Hi-C data were aligned to the Purge Haplotigs draft assembly to create a new 342 Mb assembly composed of 288 scaffolds with an N50 of 41.1 Mb and a maximum scaffold length of 52.4 Mb (Table 2; Figure 1). Among these Hi-C scaffolds, seven chromosome-length scaffolds with a total length of 299 Mb (87% of the 342 Mb genome) corresponded directly to the seven *R. argutus* chromosomes (Table 3).

Table 1. Summary of PacBio sequencing data used to generate the Burbank Thornless (*R. ulmifolius*) genome assembly

|  | PacBio |
| --- | --- |
| Number of Bases | 32,483,465,106 |
| Number of Reads | 3,313,667 |
| N50 Read Length (bp) | 18,250 |
| Mean Read Length (bp) | 9,803 |
| GC % | 37.4% |
| Nominal coverage (367 Mb genome) | 89x |

Table 2. Summary statistics for the assembled *R. ulmifolius* Burbank Thornless genome.

| Estimated genome size (flow cytometry) | 366.75 Mb |
| --- | --- |
| Total assembly length | 341.59 Mb |
| No. of scaffolds | 288 |
| No. of chromosomes | 7 |
| Size of sequence anchored on chromosomes | 299 Mb |
| Maximum scaffold length | 52.4 Mb |
| N50 scaffold length (bp) | 41.1 Mb |
| Repeat content | 48.47% |

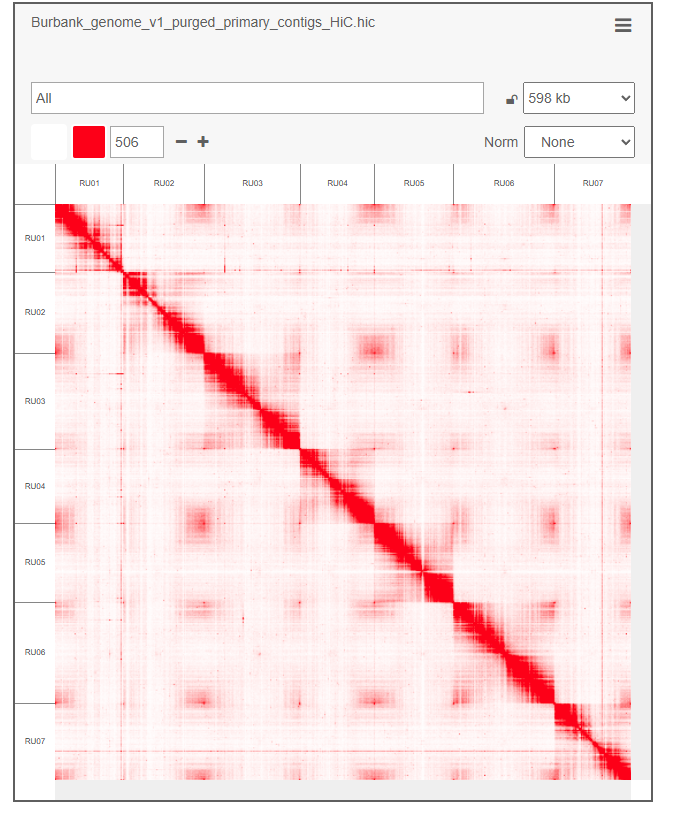

Figure 1. Hi-C interaction matrix for the ‘Burbank Thornless’ blackberry (*R. ulmifolius*) assembly. An interactive version of this map is available at https://t.3dg.io/burbank-blackberry-Fig-1 .

Table 3. Summary statistics for the seven super-scaffolds corresponding to the Burbank Thornless blackberry (*R. ulmifolius*) base chromosomes.

| Chromosomes | Total Length (bp) | N count^a^ | Gaps |
| --- | --- | --- | --- |
| Ru01 | 35232713 | 30000 | 60 |
| Ru02 | 42068357 | 26500 | 53 |
| Ru03 | 49739433 | 33500 | 67 |
| Ru04 | 38572228 | 23000 | 46 |
| Ru05 | 41135734 | 20500 | 41 |
| Ru06 | 52365428 | 28000 | 56 |
| Ru07 | 39788902 | 25000 | 50 |
| Total size (7 chromosomes) | 298902795 | 186500 | 373 |
| Unassembled fragments (281 scaffolds) | 42690562 | 21000 | 42 |
| Total genome | 341593357 | 207500 | 415 |

^a^N count refers to the total number of undetermined nucleotide bases in the assembly

***Genome size estimation***

The nuclear flow cytometry generated estimate of the *R. ulmifolius* genome size was 366.75 Mb (1C = 0.345 pg), indicating that 93.1% of the genome was incorporated in the assembly. This flow-cytometry based genome size estimate falls within the reported range of other diploid species in subgenus *Rubus* (*R. argutus*, *R. hispidus*, *R. canadensis*, *R. trivialis*, *R. canescens*, and *R. sanctus*), which was between 1C = 0.295 – 0.375 pg^8,24,25^.

**Gene prediction and annotation**

*RNA extraction, library preparation, and sequencing*

A total of 64,699,500 paired reads were generated from the four RNA-Seq libraries (two duplicate libraries of actively growing leaves and stems), with 12,215,973 to 20,119,289 paired reads per library. One SMRT cell with a library prepared from pooled RNA from the same tissue samples from leaves and stems and an additional tissue sample collected from root tips of Burbank Thornless was sequenced with Sequel II to generate a total of 6,152,928 polymerase reads with a mean length of 39,992 bp per read, an average insert length of 8,172 bp, and a mean subread length of 1,801 bp. A total of 1,254,325 non-chimeric CCS reads with a mean length of 1,187 bp were generated from these reads.

*Structural Gene Annotation*

Before structural annotation, 165.5 Mb (48.5%) of the Burbank Thornless genome was repeat masked. The final set of predicted genes contained 29,708 coding genes and, with counting alternative isoforms, 38,105 coding transcripts (Table 4). The 29,708 coding genes had an average length of 3,405 bp. Of these coding genes, 22,871 had no alternative isoforms, 4,439 had two isoforms, and 1768 had three or more isoforms (Figure 2). In the predicted set of genes, 2,179 (93.7%) complete *R. ulmifolius* genes orthologous to the eudicots_odb10 BUSCO families were identified, along with 58 (2.5%) genes with partial match. A small fraction of the BUSCO families (3.8%) were not identified among the predicted *R. ulmifolius* genes. These results suggest that the Burbank Thornless assembly and the gene complement are 96.2% complete. Evidence support for the predicted gene models was also strong, with the majority of loci supported by EST alignments and peptide homology, and over 94% achieving Cscore values above 0.9 (Table 5).

Table 4. Summary of transcript and gene model statistics for the annotated *R. ulmifolius* Burbank Thornless genome.

| Total transcripts | 38,105 |
| --- | --- |
| Primary transcripts (loci) | 29,078 |
| Alternate transcripts | 9,027 |
| Average number of exons per gene | 5.4 |
| Median exon length (bp) | 157 |
| Median intron length (bp) | 156 |
| Complete genes | 28,644 |
| Incomplete genes (missing start codon) | 275 |
| Incomplete genes (missing stop codon) | 97 |

Table 5. Evidence support for predicted gene models.

| Support type | Models evaluated | 100% | ≥95% | ≥90% | ≥75% | ≥50% |
| --- | --- | --- | --- | --- | --- | --- |
| EST overlap^a^ | 22,417 | 19,258 | 19,692 | 19,855 | 20,246 | 20,909 |
| Peptide homology^b^ | 29,078 | 3,673 | 19,416 | 21,867 | 25,224 | 27,328 |
| Cscore^b^ | 29,078 | 25,925 | 26,982 | 27,385 | 27,947 | 28,293 |

^a^EST support was evaluated for 22,417 loci with non-conflicting EST alignments in the CDS.

^b^Peptide homology and Cscore were evaluated across all 29,078 loci.

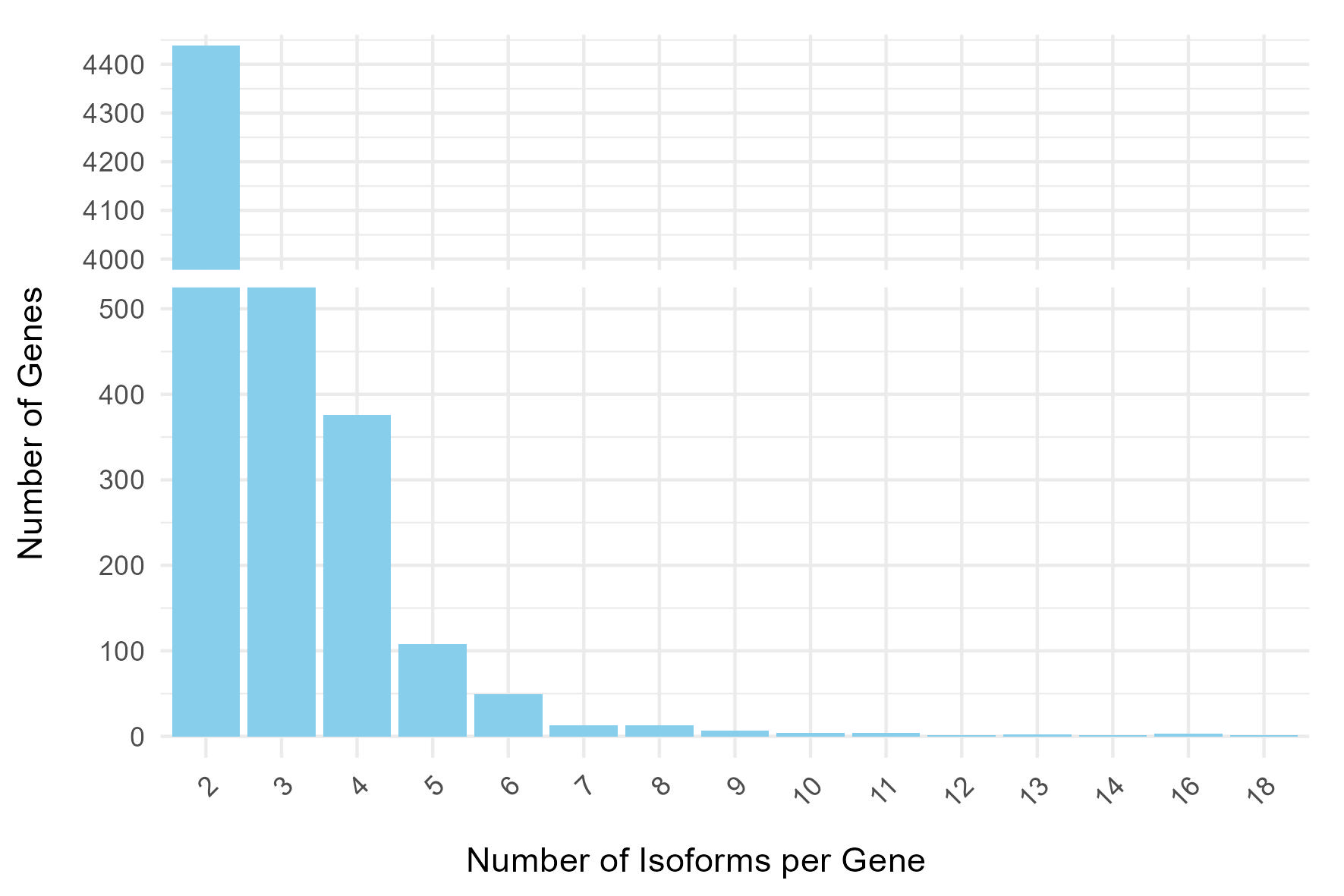

Figure 2. Distribution of the number of multiple alternative isoforms per protein-coding locus. There are 22,871 genes without alternative isoforms.

*Functional gene annotation*

Of the 38,105 predicted transcripts in the Burbank Thornless genome, a total of 37,460 (98.31%), 35,557 (93.31%), 37,312 (97.92%), 32,097 (84.23%), and 37,101 (97.37%) returned at least one hit after the blastp analysis with nr, Araport11, RefSeq, SwissProt and TrEMBL databases as subjects, respectively (Supplemental Table 4). Moreover, functional annotation analyses assigned InterPro domain, GO, KEGG pathway, and KEGG ortholog terms to 33,848 (88.83%), 25,538 (67.02%), 11,763 (30.87%) and 11,041 (28.98%) of the ‘Burbank Thornless’ predicted transcripts, respectively (Supplemental Table 5).

***Comparisons with R. argutus cv. Hillquist***

The Burbank Thornless assembly showed a high degree of collinearity with the *R. argutus* cv. Hillquist genome, with extensive alignment of nucleotide sequences across homologous chromosomes (Figure 3) and well-preserved gene order and orientation (Figure S4). The *R. argutus* genome was previously shown to have strong collinearity with *R. idaeus* cv. Anitra and *R. chingii*, with no large-scale rearrangements, translocations, or inversions observed across any of the seven chromosomes^8^. Together, these results indicate that our assembly is structurally consistent with previously published *Rubus* genomes and highlight the remarkable conservation of genome structure across the *Rubus* genus, despite divergence at the species level. Such conservation reflects the relatively recent radiation of *Rubus* species^26^ and supports the use of comparative genomics for candidate gene discovery and trait analysis within the genus.

The *R. ulmifolius* genome contained 29,708 predicted protein-coding genes, which is fewer than other published diploid *Rubus* genomes, including *R. argutus*^8^ (38,503), *R. idaeus*^27^ (39,448), *R. chingii*^28^ (33,130), and *R. occidentalis*^29^ (34,545). Despite this, our BUSCO completeness score was higher than that of *R. argutus*, and we observed a markedly higher percentage of genes with functional annotations, suggesting that our gene model set is more accurate and less inflated by fragmented or spurious predictions.

.
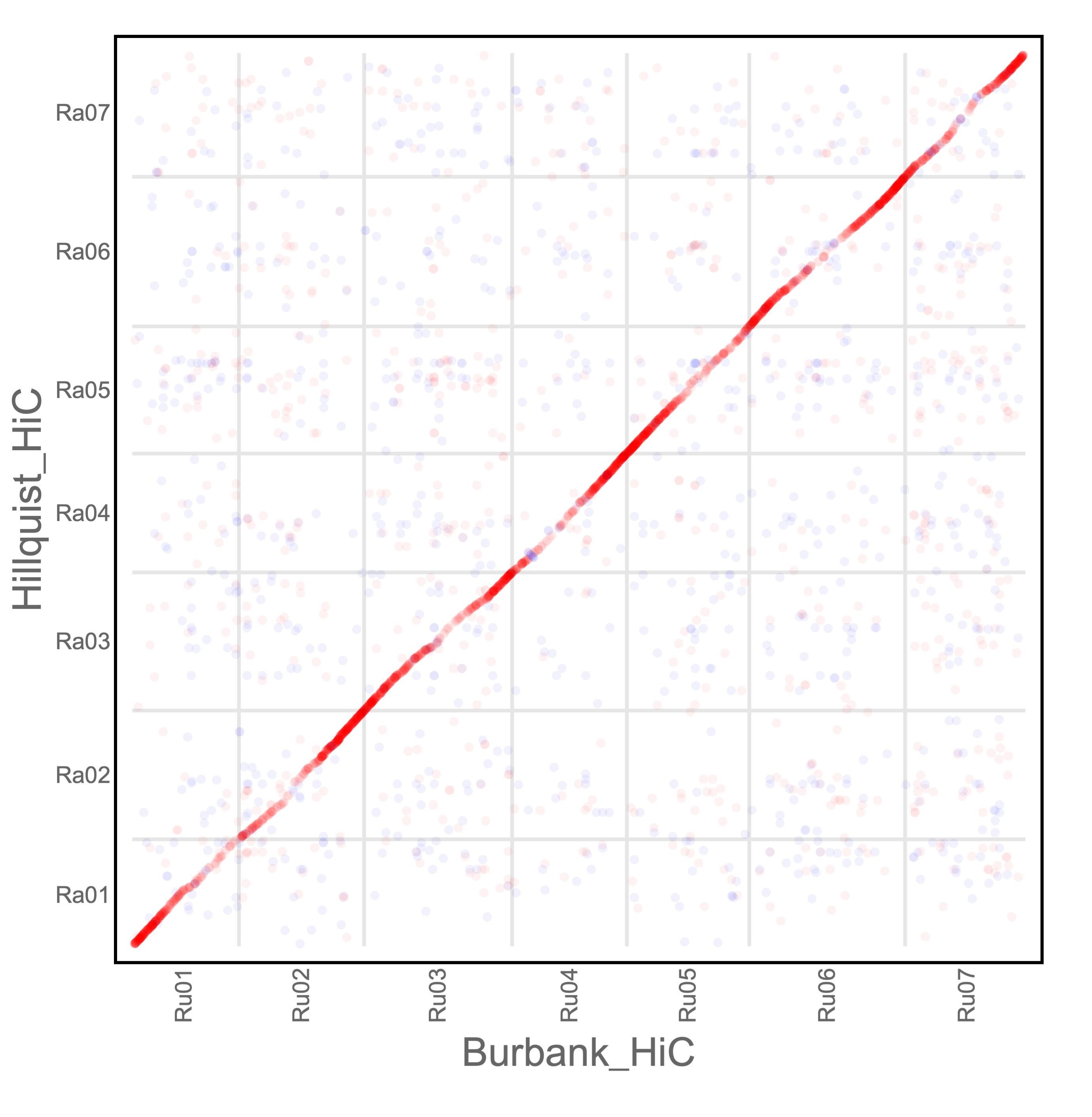

Figure 3. Macrosyteny between Burbank Thornless blackberry (*R. ulmifolius*) and Hillquist blackberry (*R. argutus*) illustrated though whole-genome alignment.

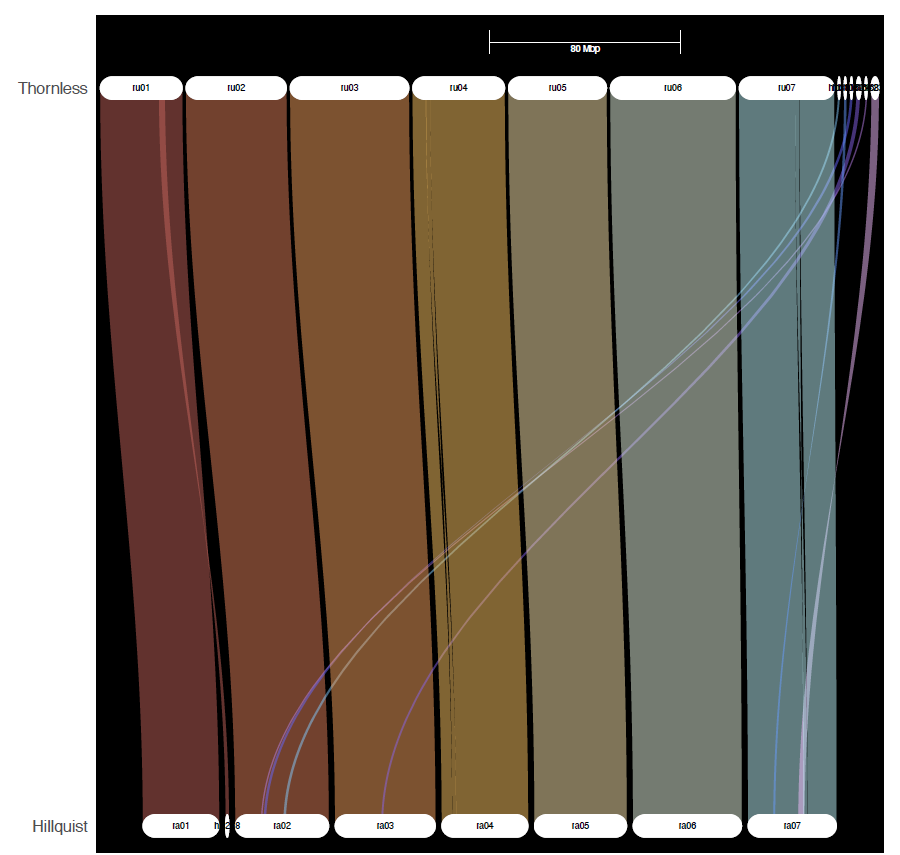

Figure 4. Macrosyteny between Burbank Thornless blackberry (*R. ulmifolius*) and Hillquist blackberry (*R. argutus*) illustrated by comparison of gene space with gene models plotted along chromosomes scaled by physical position.

**Abbreviations**

BLAST: Basic Local Alignment Search Tool; bp: base pairs; BUSCO: Benchmarking Universal Single-Copy Orthologs; BWA: Burrows-Wheeler Aligner; CCS: circular consensus sequence; cDNA: complementary DNA; CTAB: cetyl trimethylammonium bromide; Gb: gigabase pairs; GC: guanine-cytosine; GDR: Genome Database for Rosaceae; GO: gene ontology; Hi-C: high-throughput chromosome conformation capture; kb: kilobase pairs; Mb: megabase pairs; NCBI: National Center for Biotechnology Information; NCGR: National Clonal Germplasm Repository; NCSU: North Carolina State University; ORF: open reading frame; PacBio: Pacific Biosciences; RNA-Seq: RNA-sequencing; SMRT: single molecule real-time; TAIR: The Arabidopsis Information Resource; USDA-ARS, United States Department of Agriculture Agricultural Research Service.
